# Gradual cerebral hypoperfusion in a knock-in mouse model of Alzheimer’s disease triggers cortical network dysfunctions

**DOI:** 10.1101/2022.10.25.513783

**Authors:** Surjeet Singh, Sean G. Lacoursiere, Jogender Mehla, Mojtaba Nazari, Robert J. Sutherland, Robert J. McDonald, Majid H. Mohajerani

**Author notes:** Corresponding author: Majid H. Mohajerani (MHM). These authors contributed equally (Surjeet Singh, Sean G. Lacoursiere, Jogender Mehla). The Jackson Laboratory, Bar Harbor, ME 04609, USA. Department of Neurological Surgery, Washington University School of Medicine, St. Louis, MO, 63110, USA.

## Abstract

Alzheimer’s disease (AD) is characterized neuropathologically by amyloid-β (Aβ) plaques and neurofibrillary tangles. Vascular pathology caused by chronic cerebral hypoperfusion (HP) is hypothesised to exacerbate AD pathology and has emerged as an increasing cause of age-related cognitive impairment. In this study we examined the effects of gradual cerebral HP on cognitive dysfunction, Aβ pathology, microgliosis, and cortical network dynamics in C57BL/6J mice and a single App knock-in mouse model of AD (*App*^*NL-G-F*^). We performed unilateral common carotid artery gradual occlusion (UCAgO) in two-month-old mice using an ameroid constrictor. At 4 months of age, animals were tested in a behavioral battery consisting of tests of spatial learning and memory (Morris water task), recognition memory (novel object recognition task), and motor coordination (balance beam). Following behavioural testing, *in vivo* mesoscale wide-field voltage imaging was done to assess cortical functional connectivity and sensory-evoked cortical activity, and brains were harvested for pathology characterization using immunohistochemistry. We found that UCAgO reduced cerebral blood flow (CBF) in the occluded hemisphere (OH), however, subtle behavioural deficits were observed due to HP. A dissociative effect of HP was observed in resting-state functional connectivity analysis, where HP led to hyper-connectivity in C57 mice and hypo-connectivity in App mice. Interestingly, sensory stimulation of limbs contralateral to OH revealed hyper-cortical activations in the non-occluded hemisphere of C57 HP mice, however, hypo-cortical activations were observed in App HP mice. Furthermore, we found that the UCAgO increased cortical and hippocampal microgliosis in both hemispheres of C57 and App mice, a bilateral increase in Aβ deposition was only observed in App mice. These results suggest that gradual cerebral HP leads to cortical network alterations in AD, which is partly mediated via activation of microglia.

## Introduction

Alzheimer’s disease (AD) is a neurodegenerative disorder associated with extracellular amyloid-beta (Aβ) deposition within the brain parenchyma and the aggregation of the microtubule protein tau in neurofibrillary tangles in neurons (Hardy and Selkoe, 2002; Forner et al., 2017; Heneka et al., 2018). The amyloid cascade hypothesis has dominated AD research in the past few decades. Recent studies suggest that the vascular system is also a major contributor to disease progression. Vascular dysfunction and reduced cerebral blood flow (CBF) may occur prior to the accumulation and aggregation of Aβ plaques and hyperphosphorylated tau tangles (Meyer et al., 2000; de la Torre, 2002a, b, 2021). Autopsy findings in patients with dementia reveal that AD with cerebrovascular disease (mixed dementia), is more common than the ‘pure’ conditions of AD and vascular cognitive impairment (Snowdon et al., 1997; Esiri et al., 1999; Gold et al., 2007; Schneider et al., 2007; Launer et al., 2008; Schneider et al., 2009; Gorelick et al., 2011; Mazza et al., 2011; Kalaria et al., 2012; Toledo et al., 2013; Attems and Jellinger, 2014; Hattori et al., 2016; Dichgans and Leys, 2017; Feng et al., 2018; Girouard and Munter, 2018; Hartmann et al., 2018; Smith, 2018).

Large/small cerebral vasculature damage and vascular risk factors (e.g., hypertension, diabetes mellitus, atherosclerosis, smoking, hypercholesterolemia, homocysteinemia obesity) could cause cerebral hypoperfusion (HP) (McDonald, 2002; McDonald et al., 2010; Attems and Jellinger, 2014; Gardener et al., 2015; Daulatzai, 2017; van Veluw et al., 2017; Hartmann et al., 2018; Iadecola et al., 2019). However, the role of chronic cerebral HP in the pathogenesis of AD and cognitive dysfunction is unclear. Recent studies in human cohorts have presented conflicting evidence on the role of cerebral hypoperfusion in triggering AD pathology and altering brain network connectivity (Mattsson et al., 2014; Hansson et al., 2018; Fazlollahi et al., 2020; Ahmadi et al., 2021; He et al., 2021). Understanding the functional and pathogenic synergy between neurons, glia, and vascular cells could provide mechanistic insight into how alterations in cerebral blood vessels exacerbate neuronal dysfunction and underlying cognitive impairment (Iadecola, 2010; Quaegebeur et al., 2011; Zlokovic, 2011). In animal models of chronic cerebral HP, increased levels of Aβ oligomer creation/accumulation (Feng et al., 2018), pro-inflammatory cytokines (Yoshizaki et al., 2008), reduced levels of acetylcholine synthesis (Mehla et al., 2018a) and alteration of amyloid precursor protein (APP) cleavage metabolism (Bennett et al., 2000) has been observed. Further, chronic cerebral HP is shown to have negative effects on various cognitive functions, including learning and memory (Bennett et al., 1998; Kitagawa et al., 2005; Miki et al., 2009; Wang et al., 2016; Zhai et al., 2016; Feng et al., 2018). Preclinical animal models provide us an opportunity to study the contribution of vascular alterations to AD pathology (Washida et al., 2019). However, chronic cerebral hypoperfusion in animal models is mostly established by ligating the common carotid arteries leading to immediate reduction in blood flow (Farkas et al., 2006; Yoshizaki et al., 2008; Cechetti et al., 2012; Back et al., 2017; Park et al., 2019), this is a major issue, as under clinical conditions common carotid arteries are not ligated and the reduction in blood flow is also gradual.

To the best of our knowledge, no single study has evaluated the effect of unilateral common carotid gradual artery occlusion (UCAgO) on resting-state cortical connectivity, sensory-evoked cortical activity, microgliosis, amyloid pathology, and cognition in C57BL/6J and single App knock-in mouse model of AD (*App*^*NL-G-F*^) (Saito et al., 2014). An advantage of *App*^*NL-G-F*^ mouse model over other transgenic AD models is that it lacks App overexpression and toxicity and shows appreciable plaque expression and cognitive decline at six months, with clear cognitive impairment at twelve months of age (Saito et al., 2014; Saito et al., 2016; Sasaguri et al., 2017; Mehla et al., 2019). Another important aspect of our experimental design is the use of an ameroid constrictor (AC) for gradual reduction of CBF, which replicates “chronic” cerebral HP apparent in vascular cognitive impairment (Hattori et al., 2014; Hattori et al., 2015; Hattori et al., 2016). We then assessed memory and cognitive functions using the Morris water task (MWT) and novel object recognition (NOR) task (Mehla et al., 2019). Later, using *in vivo* mesoscale wide-field voltage imaging (Mohajerani et al., 2010; Mohajerani et al., 2011; Mohajerani et al., 2013; Lim et al., 2014; Chan et al., 2015), we identified resting-state functional connectivity and evoked activity pattern changes associated with HP in both C57BL/6J and *App*^*NL-G-F*^ mice (we refer to these mice as C57 and App in the remaining text). We found that UCAgO significantly reduced CBF in the occluded hemisphere (OH) and increased microgliosis in both occluded and non-occluded hemispheres (OH and NoH) of C57 and App mice. Further, UCAgO caused an increase of Aβ plaque aggregation in both hemispheres of App mice. These pathological changes resulted in mild memory impairments and alterations in functional cortical connectivity. In C57 mice, an increase in resting-state functional connectivity due to HP (hyper-connectivity) was found, whereas in App mice, functional connectivity was reduced due to HP (hypo-connectivity). Sensory stimulation of limbs contralateral to OH revealed hyper-cortical activations in non-occluded hemisphere of C57 HP mice, however, hypo-cortical activations were observed in App HP mice.

## Results

### UCAgO causes significant reduction in Cerebral Blood Flow of the occluded hemisphere in both C57 and App mice

Hypoperfusion was induced by implanting an ameroid constrictor on the left common carotid artery. We then measured CBF before UCAgO and at intervals of 1, 3, 7, 14 and 28 days following UCAgO surgery using laser speckle imaging (Mohajerani et al., 2011) to determine if the implanted ameroid constrictor would reduce CBF in the C57 and App mice. We found that following UCAgO surgery, the blood flow in the occluded hemisphere (OH) decreased gradually but significantly from the first day to the 28^th^ day [*F*(3.116, 37.39) = 7.916, *p* = 0.0003] as the ameroid constrictor began to swell and the diameter reduced (Hattori et al., 2014; Hattori et al., 2015; Hattori et al., 2016). By the 28^th^ day CBF in occluded hemisphere (OH) was significantly reduced compared to the Non-occluded hemisphere (NoH) [*F*(3,12) = 5.246, *p* < 0.05] and this effect was found in both the C57 (*p* < 0.05) and App mice (*p* < 0.005) mice (Fig 1C). The UCAgO gradually reduced blood flow over time to the ipsilateral side of the brain (OH) of the occluded artery, while blood flow to the contralateral side of occlusion (NoH) was not impacted. Furthermore, the UCAgO surgery reduced CBF equally in the C57 (*p* < 0.05) and App mice (*p* < 0.005).

**Figure 1.**
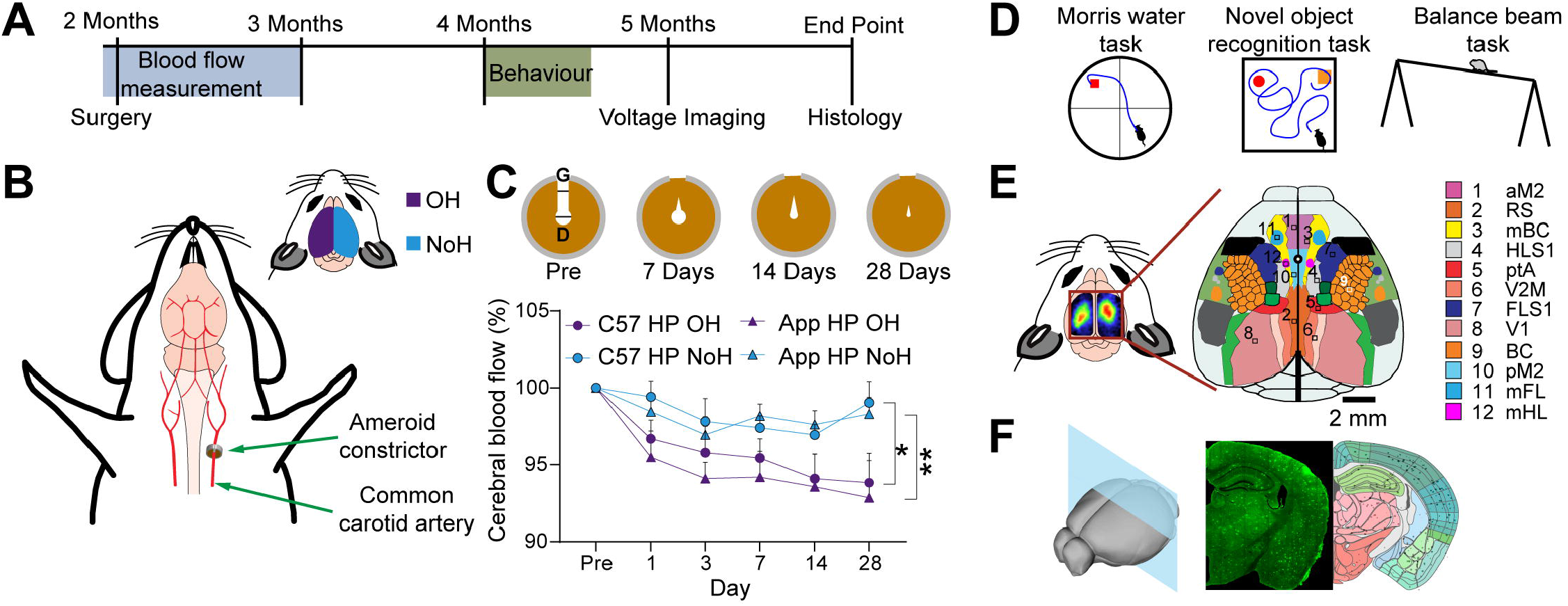
Experimental timeline. (A) Shown is a schematic illustration of the experimental design. (B) C57 and App mice underwent left common carotid artery occlusion (UCAgO) or sham surgery. The inset represents the left hemisphere as the occluded hemisphere (OH) in purple and the right hemisphere as the non-occluded hemisphere (NoH) in blue. (C) Following UCAgO surgery, the gap (g) and the internal diameter (d) of the ameroid constrictor shrank progressively and disappeared (schematic shrinkage of ameroid constrictor across 28 days based on Hattori et al., 2015). Relative CBF was measured pre-surgery and on days 1, 3, 7, 14, & 28 post-surgery using laser speckle flowmetry. The blood flow in the occluded hemisphere (OH) decreased gradually but significantly from the first day to the 28^th^ day as the ameroid constrictor began to swell and the diameter reduced (*F* (3.116, 37.39) = 7.916, *p* = 0.0003). By the 28^th^ day, CBF in the occluded hemisphere (OH) was significantly reduced compared to the Non-occluded hemisphere (NoH) (*F*(3,12) = 5.246, *p* < 0.05) in both C57 HP and App HP groups. (D) Spatial learning and memory were assessed using the Morris Water Task (MWT). Novel Object Recognition (NOR) was used to assess object learning and memory, and the Balance Beam (BB) test was performed to assess the sensory-motor function. (E) After behavioural assessment, animals were given a large bilateral craniotomy. Schematic of craniotomy showing imaged cortical regions. (F) At the experimental endpoint, mice were perfused, and tissue was collected for immunohistochemistry (IHC). \**p* < 0.05; \*\**p* < 0.01.

### Gradual cerebral HP disrupts resting-state cortical functional connectivity

After determining that UCAgO successfully reduced blood flow to the OH, we wanted to identify changes in cortical functional connectivity associated with HP in C57 and App mice. Following behavioural testing and when the mice were 5 months old, we imaged ongoing spontaneous cortical activity within both hemispheres to examine the functional connectivity using voltage sensitive dye (VSD) imaging (Mohajerani et al., 2010; Chan et al., 2015; Kyweriga and Mohajerani, 2016; Greenberg et al., 2018; Balbi et al., 2019) and calculated functional connectivity matrices based on correlation analysis. The mice were anesthetised with urethane and a 7×8 mm bilateral craniotomy (bregma 2.5 to −4.5 mm, lateral 0 to 4 mm was performed on the sham (C57, *n* = 4 and App,, *n* = 7) and HP (C57, *n* = 4 and App, *n* = 8) mice as described previously (Kyweriga and Mohajerani, 2016) and spontaneous voltage-sensitive dye (VSD) imaging of cortical responses were recorded. This method has the advantage of high spatiotemporal resolution, and large-scale recording of subthreshold and suprathreshold neuronal activity (Mohajerani et al., 2013). With these advantages in mind, we assessed mesoscale functional connectivity and plasticity after UCAgO across both hemispheres and in both C57 and App mice.

To access functional connectivity disruptions due to HP, a region-based cortical correlation analysis was performed on resting state spontaneous VSD imaging data. Twelve, 5 × 5-pixel regions of interest (ROIs) were selected from each hemisphere for a total of 24 cortical responses. Zero-lag Pearson correlation coefficient of ROI time courses was calculated to generate the functional connectivity matrix. To represent the data as a whole and to show relationships between regions within hemisphere (*intra* hemispheric) and between hemispheres (*inter* hemispheric), a 24 × 24 correlation-based functional connectivity matrix measured from resting-state spontaneous activity was created for each group (C57 Sham, C57 HP, App Sham, and App HP) (Fig S1). In the sham C57 connectivity matrix (Fig. S1A*i*), well known relationships can be identified in the connectivity matrix, such as between the somatosensory barrel cortex and motor cortex as shown previously (Mohajerani et al., 2010).

For C57 mice, the difference of mean correlation matrix (C57 HP – C57 Sham) revealed that gradual cerebral HP in the C57 mice led to increased cortical connectivity (hyperconnectivity) as represented by warmer colours (red) in fig. 2A (see Fig. S1A*i*-*ii* for mean correlation matrices).

**Figure 2.**
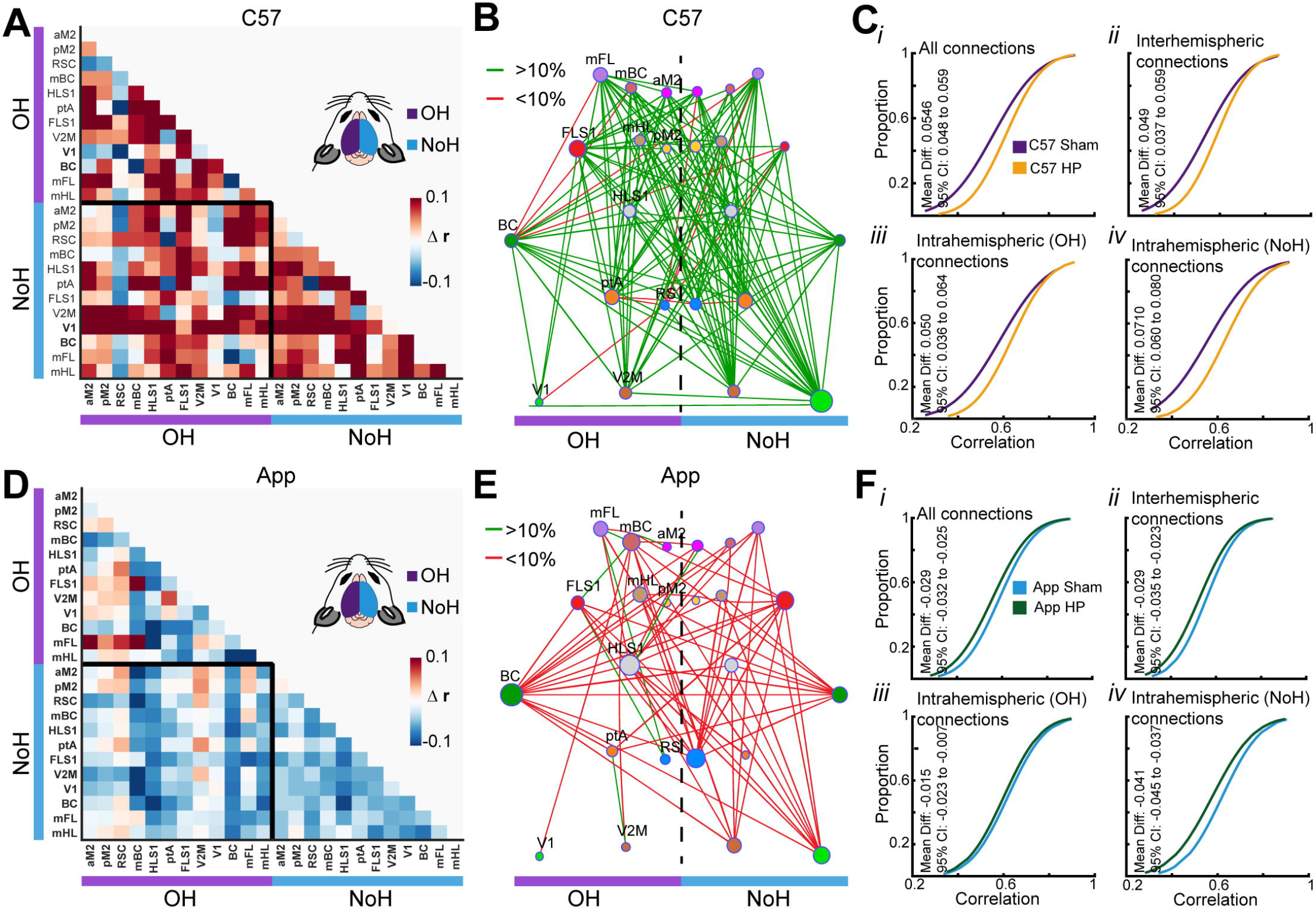
Unilateral gradual cerebral hypoperfusion resulted in dissimilar patterns of cortical functional connectivity in C57 and App mice. Cortical functional connectivity matrices were generated based on zero-lag correlation of resting-state spontaneous activity in 24 cortical regions of interest. Since this is a symmetric matrix only values from the lower triangular matrix are used for analysis. (A) Matrix showing difference in mean correlation of C57 HP and C57 Sham animals (C57 HP – C57 Sham; n = 4 mice for both groups), regions in the occluded (OH) and non-occluded (NoH) hemispheres are highlighted by a purple and blue lines respectively. The values inside the black square represents changes in inter-hemispheric connections, the upper left and the bottom right triangles represents intra hemispheric changes in OH and NoH respectively. Warmer colours (red) highlight increased cortical connectivity due to HP in C57 animals. (B) To better demonstrate the changes between C57 Sham and C57 HP animals, correlation differences were color-coded to illustrate changes in network strength. Red indicates a loss of strength (<10% change in strength), and green indicates a gain of strength (>10% change in strength). Network graph of changes in cortical connections (HP-effect) shows that HP in C57 animals leads to hyperconnectivity in cortical network. Red and green lines represent connection for which functional connectivity reduced (24 connections) or increased (278 connections) by 10% in C57 HP group compared to C57 group. (C) Cumulative distribution functions (*cdf*) of correlation values suggest increased cortical connectivity in C57 HP group as compared to C57 Sham group. These changes in cortical connectivity strength were accessed using Generalized linear mixed-effects (GLME) models (n = 4; C57 Sham and n = 4; C57 HP). (*i*) all connections: 276 connections/animal; GLME HP-effect t(2206) = 19.39, p < 0.001; (*ii*) interhemispheric connections: 144 connections/animal; GLME HP-effect *t*(1150) = 12.328, *p* < 0.001; (*iii*) intrahemispheric (OH) connections: 66 connections/animal; GLME HP-effect *t*(526) = 6.9296, *p* < 0.001; and (*iv*) intrahemispheric (NoH) connections: 66 connections/animal; GLME HP-effect *t*(526) = 17.133, *p* < 0.001. (D) Similar to A, but for App HP - App Sham (n = 8 for App HP, and n= 7 for App Sham) representing reduced cortical connectivity due to HP in App mouse model. (E, and F) Similar to B, and C but for App HP - App Sham. Cooler colours (blue) highlight reduced cortical connectivity due to HP in App animals. Red and green lines represent connection for which functional connectivity reduced (122 connections) or increased (12 connections) by 10% in App HP group compared to App group. Cumulative distribution functions (*cdf*) of correlation values reveal reduced cortical connectivity in App HP group as compared to App group. (*i*) all connections: GLME HP-effect *t*(4138) = 7.2909, *p* < 0.001; (*ii*) interhemispheric connections: GLME HP-effect *t*(2158) = 5.2929, *p* < 0.001; (*iii*) intrahemispheric (OH) connections: GLME HP-effect *t*(988) = 1.382, n.s.; and (*iv*) intrahemispheric (NoH) connections: GLME HP-effect *t*(988) = 6.1703, *p* < 0.001.

To demonstrate the effect of UCAgO on functional connectivity, we created a network graph using modified MATLAB scripts (Lim et al., 2015) and the brain connectivity toolbox (https://sites.google.com/site/bctnet/; (Rubinov and Sporns, 2010)). Connections with more than 10% change in correlation strength following HP are shown, with red and green lines representing reduced and increased functional connectivity respectively. Network graph in fig. 2B reveal a global increase in inter-regional functional connectivity in C57 HP mice (Fig. 2B) indicating hyperconnectivity.

For qualitative examination of functional connectivity changes we plotted cumulative distribution function (*cdf*) of correlation values. Changes in functional connectivity were assessed using Generalized linear mixed-effects (GLME) models. When all nodes of connectivity matrices were considered, we found a significant increase in functional connectivity in the C57 HP mice compared to the C57 sham mice [*t*(2206) = 19.39, *p* < 0.001; GLME HP-effect, all connections/mouse = 276; Fig. 2C*i*]. The same pattern of increase in functional connectivity was also found across hemispheres [*t*(1150) = 12.328, *p* < 0.001; GLME HP-effect, *inter*hemispheric connections/mouse = 144; Fig. 2C*ii*], and within both the OH hemisphere [*t*(526) = 6.9296, *p* < 0.001; GLME HP-effect, *intra*hemispheric (OH) connections/mouse = 66; Fig. 2C*iii*] and NoH [*t*(526) = 17.133, *p* < 0.001; GLME HP-effect, *intra*hemispheric (NoH) connections/mouse = 66; Fig. 2C*iv*].

Interestingly, for App mice gradual cerebral HP led to reduction in functional connectivity (hypoconnectivity) as shown by cooler colours (blue) more negative values in the difference correlation matrix (Fig. 2D; see Fig. S1B*i*-*ii* for mean correlation matrices). Further, network graph of changes in ROI correlation values revealed a global reduction in inter-regional functional connectivity in App mice (Fig. 2E). Considering all nodes of connectivity matrices, we found that App HP mice have significantly reduced connections compared to the App sham mice [*t*(4138) = 7.2909, *p* < 0.001; GLME HP-effect, all connections/mouse = 276; Fig. 2F*i*]. When looking across hemispheres the same pattern was observed [*t*(2158) = 5.2929, *p* < 0.001; GLME HP-effect, *inter*hemispheric connections/mouse = 144; Fig. 2F*ii*]. Within hemispheres, we found no significant difference in the OH [*t*(988) = 1.382, *p* = 0.167; GLME HP-effect, *intra*hemispheric (OH) connections/mouse = 66; Fig. 2F*iii*] but we did in the NoH [*t*(988) = 6.1703, *p* < 0.001; GLME HP-effect, *intra*hemispheric (NoH) connections/mouse = 66; Fig. 2Fi*v*].

Overall, we found that gradual cerebral HP increased functional connectivity (*hyper connectivity*) in C57 mice while reducing functional connectivity (*hypo connectivity*) in App mice. These differential effects of cerebral HP on functional connectivity strength were consistently observed in *inter*-, *intra-* and overall connections of C57 and App mice, even though the reduction in blood flow was confined to the OH.

### Gradual cerebral HP altered sensory-evoked cortical activity

We hypothesized that in addition to spontaneous activity alterations with HP, sensory-evoked cortical responses will also be affected. Patterns of sensory signal processing are shown to be altered in mouse models of AD and after targeted mini-strokes (Sigler et al., 2009; Mohajerani et al., 2011; Maatuf et al., 2016). To assess how patterns of sensory-evoked cortical activity changes in our experiments, forelimb (FL) and hindlimb (HL) stimulation-evoked VSD signals were recorded in both cortical hemispheres of sham and HP mice.

Alteration in population responses were compared based on the following six parameters: rise time, fall time, inter-hemispheric delay, peak amplitude (Δ*F*/*F*_0_), area under the curve (AUC) and laterality index (see Methods).

As previously described for the non-stroke conditions (Ferezou et al., 2007; Mohajerani et al., 2011) we also found that sensory stimulation of FL or HL leads to first activation in contralateral hemisphere and the signal later (∼20 ms delay) propagates to hemisphere ipsilateral to stimulated limb. In addition, the secondary response in the ipsilateral hemisphere is lower in magnitude as compared contralateral hemisphere (Fig. 3A*i*, 4A*i*, S4A*i* & S5A*i* – C57 sham).

**Figure 3.**
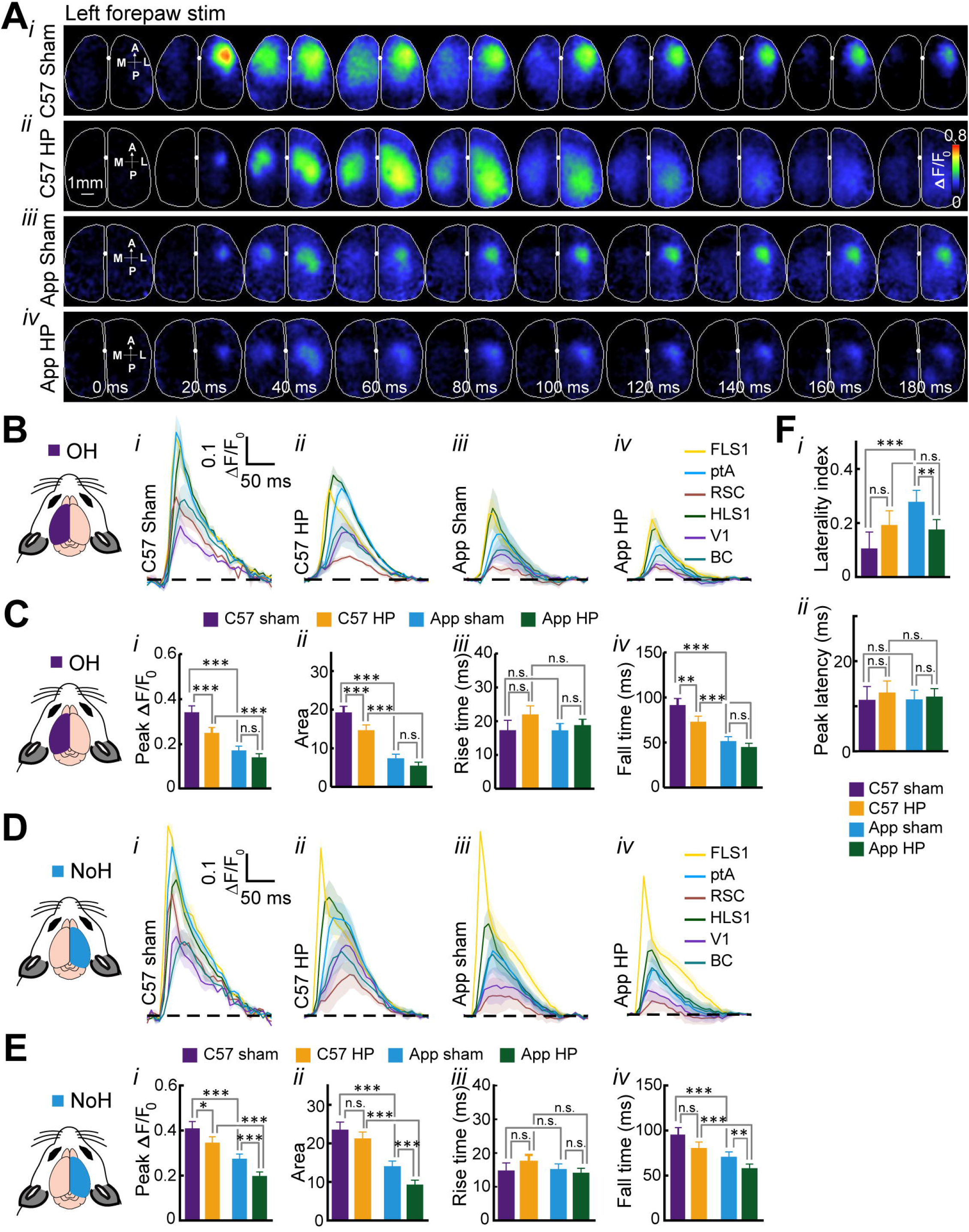
Reduced sensory evoked cortical activation after left forelimb (FL) stimulation in C57 HP and App HP mice. (A*i-iv*) Representative patterns of bilateral cortical activation following 1 mA, 1 ms pulse stimulation of left FL in C57 sham, C57 HP, App sham, and App HP mice. The VSDI montages represents 10 frames of evoked cortical responses at interval of 20 ms after stimulus onset (0.00 ms). The first image in the montage indicates the anterior (A), posterior (P), medial (M) and lateral (L) directions. (B*i-iv*, D*i-iv*) Average VSD signal from representative six regions of interest (ROIs) of OH and NoH in response to left forelimb stimulation. Primary hindlimb and forelimb sensory areas (HLS1 and FLS1), parietal associational area (ptA), retrosplenial cortex (RS), primary visual cortex (V1), barrel cortex (BC), as estimated using stereotaxic coordinates (Paxinos & Franklin, 2004). (C) In OH, there was a significant reduction of peak Δ*F*/*F*_0_ (C*i*) and area under Δ*F*/*F*_0_ – time curve (C*ii*) for C57 HP group as compared to C57 sham group (*p* < 0.0001), but no change was observed in App sham and HP groups, further no differences in rise time (C*iii*) were observed, however, fall time (C*iv*) was significantly different among C57 sham and C57 HP group (*p* < 0.01). (E) In NoH, significant reduction in peak Δ*F*/*F*_0_ (E*i*) (*p* < 0.0001), area under Δ*F*/*F*_0_ – time curve (E*ii*) (*p* < 0.0001) and fall time (E*iv*) (*p* < 0.01) was observed for App sham and App HP group, there was no difference in rise time across all groups (E*iii*). These results suggest reduced cortical activations in OH and NoH due to both HP and AD pathology for FL left stimulation. (F) Comparison of laterality index (F*i*) revealed significant reduction in App HP group compared to App sham group (*p* < 0.01), but no difference was observed in peak latency (F*ii*) for any group. App animals (n = 7 Sham; n = 7 HP) and C57 animals (n = 4 Sham; n = 4 HP). All values are expressed as mean ± SEM. * *p* < 0.05; ** *p* < 0.01; *** *p* < 0.001.

For left FL stimulation (Fig. 3B-F) we found a significant effect of HP and strain in peak amplitude reduction in both OH (HP: [*F*(1,215) = 24.76, *p* < 0.0001], strain: [*F*(1,215) = 128.49, *p* < 0.0001]) and NoH (HP: [*F*(1,215) = 28.04, *p* < 0.0001], strain: [*F*(1,215) = 113.89, *p* < 0.0001]). Similar effects were observed for area under the curve in both OH (HP: [*F*(1,215) = 21.99, *p* < 0.0001], strain: [*F*(1,215) = 230.41, *p* < 0.0001]) and NoH (HP: [*F*(1,215) = 17.82, *p* < 0.0001], strain: [*F*(1,215) = 164.20, *p* < 0.0001]). No significant differences in rise time of the evoked were observed however there was a significant effect of HP and strain on fall time of the evoked signal was observed in both OH (HP: [*F*(1,215) = 15.88, *p* < 0.0001], strain: [*F*(1,215) = 116.39, *p* < 0.0001]) and NoH (HP: [*F*(1,215) =

16.80, *p* < 0.0001], strain: [*F*(1,215) = 48.70, *p* < 0.0001]). No significant changes in peak latency were observed, but there was significant effect of strain ([*F*(1,215) = 9.45, *p* = 0.002]) and HP x strain interaction ([*F*(1,215) = 13.79, *p* < 0.001]) on laterality index. See Fig. S2 for left FL stimulation related region-specific comparisons of rise time, fall time, inter-hemispheric delay, peak amplitude (Δ*F*/*F*_0_), area under the curve (AUC) and laterality index. Overall, these results suggest reduced cortical activations due to both HP and AD for FL left stimulation.

Further, for right FL stimulation (Fig. 4B-F) the results for OH were similar to left FL stimulation-OH, however, we observed some unique changes in NoH for right FL stimulation. Interestingly, there was no effect of HP on peak amplitude or area under the curve in NoH, however there was significant HP x strain interaction for both peak amplitude [*F*(1,215) = 5.57, *p* = 0.019] and area under the curve [*F*(1,215) = 12.44, *p* < 0.001]. We observed that C57 HP mice had higher activation in NoH as compared to C57 Sham, although this was not statistically significant. On further evaluation, a significant effect of HP [*F*(1,215) = 36.00, *p* < 0.0001], strain [*F*(1,215) = 20.14, *p* < 0.001] and HP x strain interaction [*F*(1,215) = 23.78, *p* < 0.001] was observed on changes in laterality index. Interestingly, for C57 HP mice, right FL stimulation not only led to reduction of cortical activation in OH but also increased NoH cortical activation as suggested by negative laterality index values. See Fig. S3 for right FL stimulation related region-specific comparisons of rise time, fall time, inter-hemispheric delay, peak amplitude (ΔF/F0), area under the curve (AUC) and laterality index.

**Figure 4.**
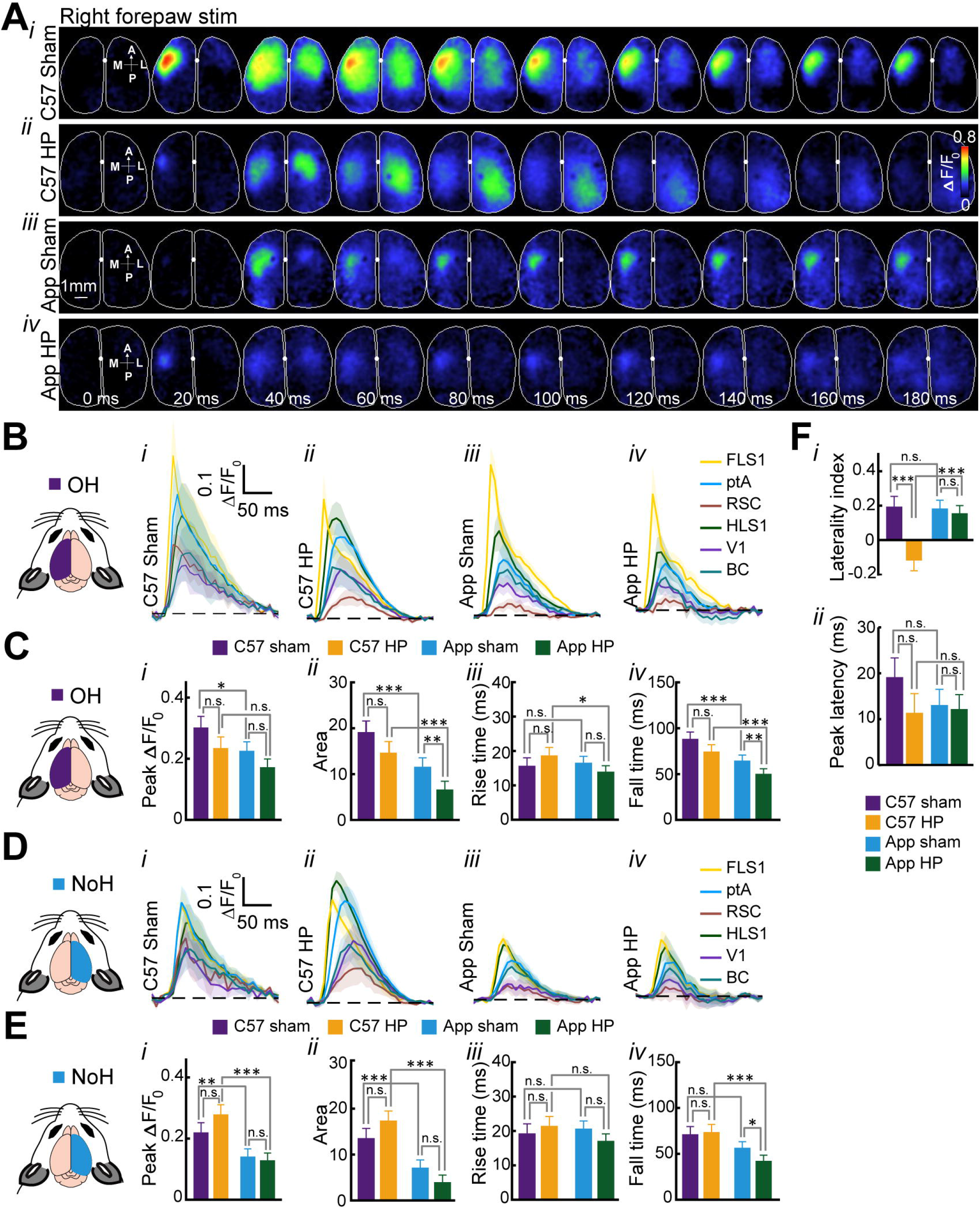
Increased sensory evoked cortical activation in non-occluded hemisphere (NoH) of C57 HP mice after right forelimb (FL) stimulation. (A*i-iv*) Representative patterns of bilateral cortical activation following 1 mA, 1 ms pulse stimulation of right FL in C57 sham, C57 HP, App sham, and App HP mice. The VSDI montages represents 10 frames of evoked cortical responses at interval of 20 ms after stimulus onset (0.00 ms). The first image in the montage indicates the anterior (A), posterior (P), medial (M) and lateral (L) directions. (B*i-iv*,D*i-iv*) Average VSD signal from representative six regions of interest (ROIs) of OH and NoH in response to right forelimb stimulation. Primary hindlimb and forelimb sensory areas (HLS1 and FLS1), parietal associational area (ptA), retrosplenial cortex (RS), primary visual cortex (V1), barrel cortex (BC), as estimated using stereotaxic coordinates (Paxinos & Franklin, 2004). (C) In OH, there was a significant reduction in area under Δ*F*/*F*_0_ – time curve (C*ii*) and fall time (C*iv*) for App HP group as compared to App sham group (*p* < 0.01), but no change is observed in C57 sham and C57 groups, further no differences in peak Δ*F*/*F*_0_ (C*i*) and rise time (C*iii*) were observed for C57/App sham and HP groups. (E) In NoH, no significant change was observed in peak Δ*F*/*F*_0_ (E*i*), area under Δ*F*/*F*_0_ – time curve (E*ii*) and rise time (E*iii*) across C57/App sham and HP groups, however, there was significant decrease in fall time (E*iv*) of App HP group as compared to App sham group (*p* < 0.05). These results suggest reduced cortical activations in OH and NoH due to both HP and AD pathology for FL right stimulation. (F) Significant negative laterality index (F*i*) for C57 HP mice (*p* < 0.0001) suggest that there is not only reduction of cortical activation in OH but also increase in NoH cortical activation, no significant difference was observed in peak latency (F*ii*) for any group. App animals (n = 7 Sham; n = 8 HP) and C57 animals (n = 4 Sham; n = 4 HP). All values are expressed as mean ± SEM. * *p* < 0.05; ** *p* < 0.01; *** *p* < 0.001.

Interestingly, for left and right HL stimulus (Fig. S4B-F & S5B-F) we observed changes similar to left and right FL stimulus. See Fig. S6 & S7 for left and right HL stimulation related region-specific comparisons of rise time, fall time, inter-hemispheric delay, peak amplitude (ΔF/F0), area under the curve (AUC) and laterality index.

An important finding from these sensory evoked experiments is that stimulation of limbs contralateral to OH leads to an increased response in the NoH for C57-HP mice, also shown by the negative laterality index in such stimulations. These results suggest that for C57-HP mice, NoH might be compensating for the HP in OH, however, for App-HP such a compensatory mechanism may not be plausible, probably due to underlying AD pathology.

### Gradual cerebral HP increased microgliosis and Aβ plaque throughout the brain

We quantified 82e1 and Iba1 staining in the OH and NoH of the cortex and HPC of C57 and App mice to investigate the effects of gradual cerebral HP on Aβ plaque and microgliosis, respectively. Despite the longitudinal differences in cerebral blood flow in the OH and NoH over 28 days following constrictor implantation, we found no hemispheric differences in cortical microgliosis [*F*(1, 26) = 0.0055, *p* = 0.941] or Aβ plaque [*F*(1, 26) = 0.00197, *p* = 0.965]. Similar trend was observed for HPC microgliosis [*F*(1, 26) = 0.1138, *p* = 0.739] and Aβ plaque [*F*(1, 26) = 0.0150, *p* = 0.904] (Fig. S8A-B). Therefore, the data from both hemispheres was combined.

Gradual cerebral HP significantly increased microgliosis in the cortex [*F*(1, 30) = 23.20, *p* < 0.0001; two-way ANOVA; Fig. 5B*i*] and HPC of both App and C57 mice [*F*(1, 30) = 37.42, *p* < 0.0001; Fig. 5B*ii*]. The cortex of the App mice had significantly greater microgliosis compared to C57 mice [*F*(1, 30) = 13.67, *p* = 0.0009] but similar levels of microgliosis in the HPC [*F*(1, 30) = 2.31, *p* = 0.1387]. The effect of HP in the cortex was significantly greater in App mice compared to C57 mice as evidenced by a significant strain x HP interaction [*F*(1, 30) = 11.60, *p* = 0.0019] but this interaction was not found in the HPC [*F*(1, 30) = 1.914, *p* = 0.177].

**Figure 5.**
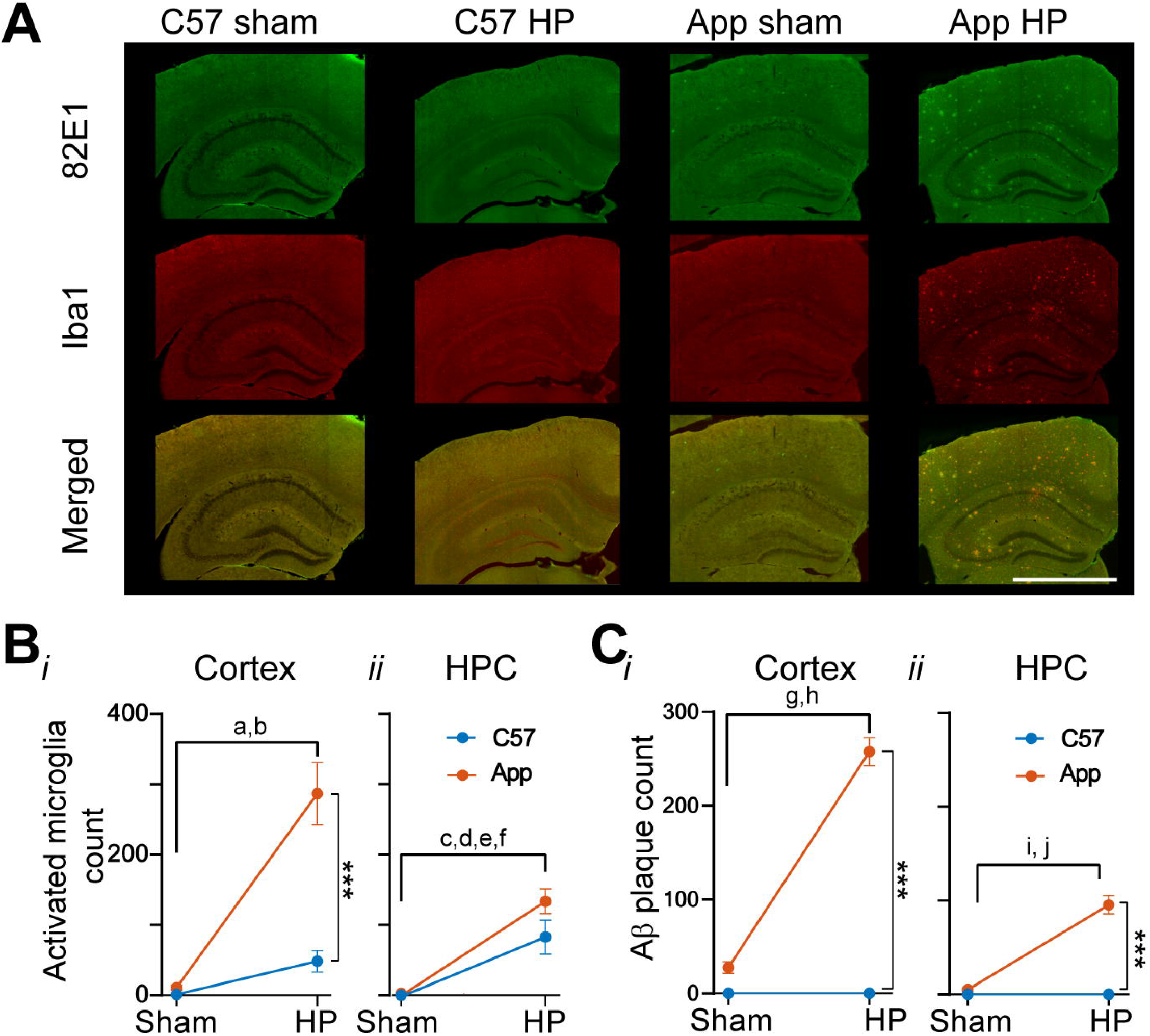
Gradual cerebral HP was found to significantly exacerbate microgliosis and Aβ pathology in the App mice. (A) Representative photomicrographs of immunohistochemistry staining of microgliosis (Iba1), Aβ plaque (82E1), and combined (82E1 + Iba1), images of cortex and HPC of the C57 sham, C57 HP, App sham, and App HP mice. Scale bar for Iba1, 82E1, and combined is 1250 μm. (B) Activated microglia count in cortex (*i*) and HPC (*ii*) was significantly increased in the cortex of the App HP mice (*n* = 6) compared to the App mice (*n* = 5; *p* < 0.005). Furthermore, the cortex of the App HP mice showed significantly greater microglia count compared to the C57 HP mice (*n* = 5; *p* < 0.05) and C57 mice (*n* = 5; *p* < 0.001). The HPC in the App HP mice showed increased microglial count compared to the App mice (*p* < 0.005). (C) Aβ plaque count in both cortex (*i*) and HPC (*ii*) was significantly greater in the App HP mice compared to the App sham mice. All values are expressed as mean ± SEM. * *p* < 0.05; ** *p* < 0.01; *** *p* < 0.001.

HP significantly increased Aβ plaque deposition in both the cortex [*F*(1, 30) = 105.0, *p* < 0.0001; Fig. 5C*i*] and HPC [*F*(1, 30) = 47.66, *p* < 0.0001; Fig. 5C*ii*] of App mice. No plaque was found in C57 Sham or HP mice.

Gradual cerebral HP
significantly increased microgliosis and Aβ plaque in the cortex and HPC of App mice. The effect of HP in the cortex on microgliosis and Aβ deposition was significantly greater in App mice compared to C57 mice, but the effect on microgliosis was observed in the HPC as microgliosis increased to similar levels in both C57 and App mice.

### Behavioural changes associated with HP in spatial learning, fine sensory motor abilities, object memory

Together with changes in functional cortical connectivity/activity and AD-like pathology we wanted to determine if gradual cerebral HP impaired behaviour.

We used the MWM to test spatial memory using the protocol previously described in (Mehla et al., 2019). Briefly, the mice were trained to find a submerged platform over 8 days followed by a no-platform probe trial on the 9^th^ day. We found that HP had no effect on escape latency [*F*(1, 42) = 2.040, *p* = 0.161; Fig. 6A]. When the mice were grouped into C57 and App groups regardless of HP condition, C57 and App mice significantly reduced their escape latency across training [*F*(7, 308) = 29.76, *p* < 0.0001] but C57 mice found the platform quicker [*F*(1, 44) = 76.19, *p* < 0.0001] and reduced their escape latency to a greater extent [*F*(7, 308) = 2.582, *p* = 0.0134].

**Figure 6.**
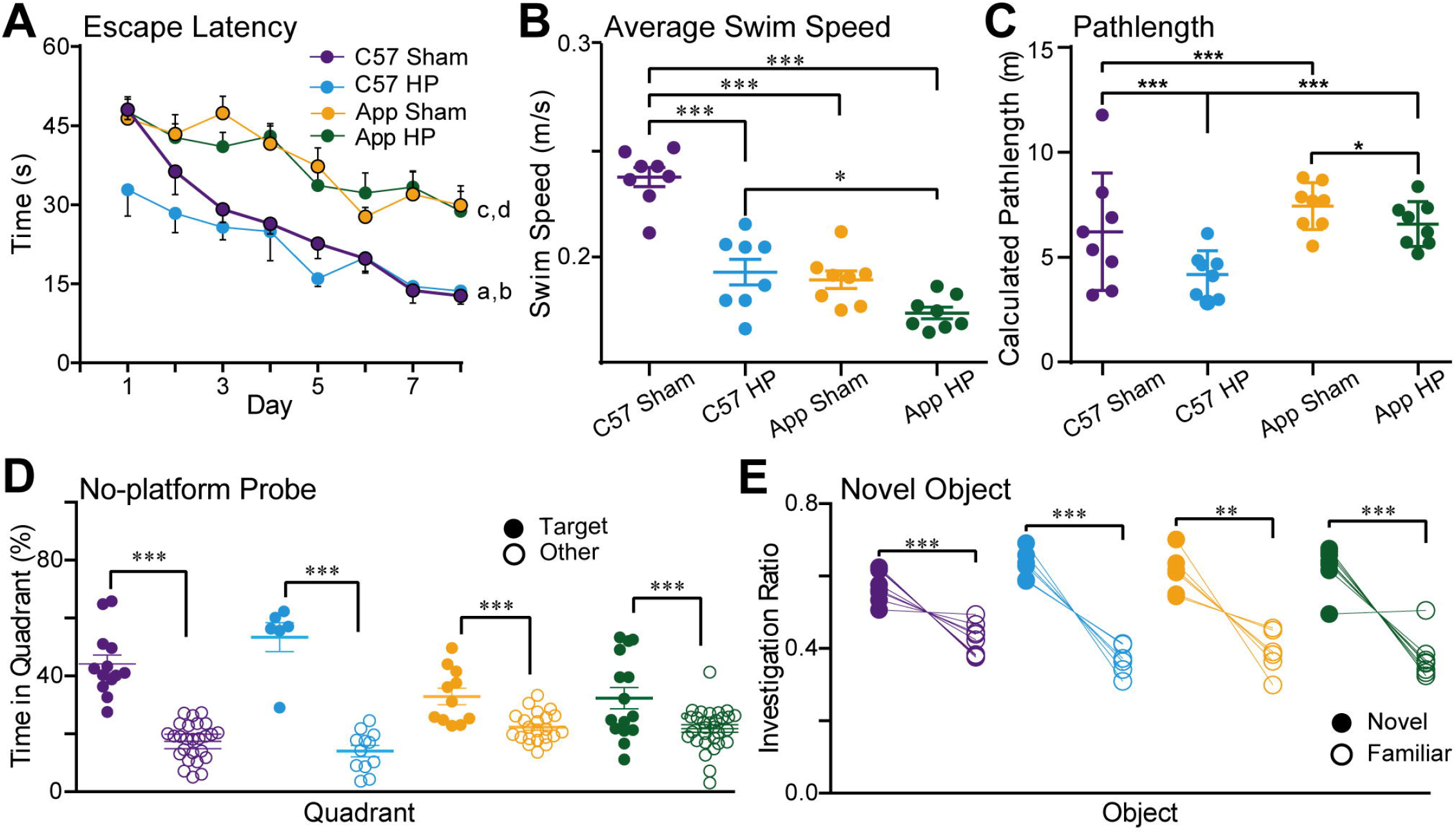
Gradual cerebral HP did not impair spatial memory or fine sensory motor abilities or object recognition. (A-C) Spatial learning and memory performance in the MWT: (A) we found that all mice significantly shortened their latency to find the hidden platform target from day 1 to day 8; a. C57 sham, *p <* 0.0001; b. C57 HP, *p* < 0.01; c. App sham, *p* < 0.05; d. App HP, *p* < 0.001. The C57 mice had significantly shorter latency overall compared to the App mice (*p* < 0.0001). (B) The C57 sham mice swim speed was significantly faster than the C57 HP, App sham, and App HP (*p* < 0.0001) mice. The App HP mice have an average swim speed significantly slower than C57 HP mice (*p* < 0.05). (C) A path length analysis showed that C57 mice had a significantly shorter average path length compared to the App mice and HP lead to further reduction in the path length. (D) The no-platform probe trial showed that all mice spent a significantly more time in the target quadrant compared to the non-target quadrants. The C57 mice spent a significantly higher percentage of time in the target quadrant compared to the App mice (*p <* 0.05). No difference was found in target preference percent between the C57 sham and C57 HP mice, nor between the App sham and App HP mice. €The NOR task showed the mice spent significantly more time investigating the novel object compared to the familiar object. All values are expressed as mean ± SEM. * *p* < 0.05; ** *p* < 0.01; *** *p* < 0.001.

Swim speed was significantly reduced by HP [*F*(1, 42) = 11.04, *p* = 0.0019; Fig. 6B] but on average C57 mice swam significantly faster than the App mice [*F*(1, 42) = 13.80, *p* = 0.0006]. Using a Sidak’s multiple comparison, we found that C57 Sham mice swam significantly faster than all groups of mice (*p* < 0.05) but we found that App Sham and App HP mice had similar swim speeds (*p* > 0.05); however, a t-test reveals that the difference in swim speed between App Sham and HP mice is significantly different [*t*(14) = 3.13, *p* = 0.007].

A path length analysis revealed that the C57 mice on average had a shorter pathlength compared to App mice [*F*(1, 42) = 39.11, *p* < 0.0001]. Gradual cerebral HP resulted in a significantly shorter path length in both C57 and App mice [*F*(1, 42) = 24.86, *p* < 0.0001; Fig. 6C] but with a greater reduction in C57 mice [*F*(1, 42) = 4.127, *p* = 0.049].

Despite a decrease in both average speed and path length due to HP, distance by speed ratio was conserved for both sham and HP group which would explain why latency to find the platform was similar between control and HP mice (Fig. 6A-C).

Following training, a no-platform probe trail was done to determine if the mice learned the platform location. The percent time spent in the target quadrant was compared to the percent time spent in the other, non-target quadrants (Fig. 6D). One mouse was an identified outlier (ROUT Q = 1%) and removed from analysis. The mice spent significantly more time in the target quadrant compared to the other quadrants [F(1, 83) = 110.0, *p* < 0.0001]. HP was found to have no significant effect on this memory task [*F*(1, 83) = 0.512, *p* = 0.476]. But overall, C57 mice spent significantly more time in the target quadrant compared to App mice [*F*(1, 83) = 6.657, *p* = 0.012]. These results suggest that HP had no significant effect on spatial learning and memory, but HP did impair swim speed.

The novel object recognition task was completed to test object memory. As mice prefer novelty, it is predicted they will spend more time investigating the novel object compared to the already familiar object and that the combined effects of gradual cerebral HP and Aβ pathology would impair this ability. Overall, we found that the mice investigated the novel object significantly more than the familiar object [*F*(1, 35) = 63.30, *p* < 0.0001, Fig. 6E] with C57 mice investigating the novel object significantly more than the App mice [*F*(1, 35) = 117.0, *p* < 0.0001]. We found that in comparison to the sham mice, C57-HP and App-HP mice spent more time investigating the novel object than the familiar object [*F*(1, 35) = 468.2, *p* < 0.0001]. The balance beam was used to assess sensorimotor function (Mehla et al., 2018a). We found no strain [*F*(1, 33) = 0.090, *p* = 0.766] or HP [*F*(1, 33) = 0.501, *p* = 0.484] effects on time taken to cross the beam (data not shown).

## Discussion

It is hypothesized that chronic cerebral hypoperfusion can trigger the chain of pathological events in AD (Zlokovic, 2011; Pimentel-Coelho et al., 2013; Binnewijzend et al., 2016; Iturria-Medina et al., 2016; Park et al., 2019), however, some recent clinical studies contradict this hypothesis suggesting cerebral hypoperfusion to be a non-causal event or a late pathological event in the course of AD (Hansson et al., 2018; Ahmadi et al., 2021). In animal models, cerebral hypoperfusion has been shown to trigger AD related pathologies (e.g. increase in Aβ plaques, microgliosis) (Farkas et al., 2006; Yoshizaki et al., 2008; Yamada et al., 2011; Cechetti et al., 2012; Back et al., 2017; Park et al., 2019), it is important to note that chronic cerebral hypoperfusion in these studies is mostly established by ligating the common carotid arteries, this poses a major issue for the interpretation of the results, as under clinical conditions common carotid arteries are not ligated, instead there is gradual occlusion. The present study aimed at investigating if gradual cerebral hypoperfusion could induce or exacerbate AD related pathologies in C57 and App mice, and whether these changes in blood flow and pathology could be detrimental to cortical activity (spontaneous/evoked) and cognition as observed through VSD imaging and behavioural testing.

We found that UCAgO significantly reduced blood flow to the ipsilateral side of the occlusion in both C57 and App mice. Despite the reduction of blood flow, we did not find any hemisphere specific alterations in AD related disease pathology. Instead, we found that there is an overall increase in Aβ plaque and microglia count in App HP mice and an increase in microglia count in C57 HP mice. Further, no significant memory deficits were observed in MWT, although there was a reduction in swim speed due to HP. Gradual cerebral HP has been shown to increase Aβ pathology and microgliosis in mouse models of AD (Shang et al., 2016; Zhai et al., 2016; Feng et al., 2018; Shang et al., 2019) however, this increase in disease related pathology has subtle to no effect on behaviour (Hattori et al., 2015; Zhai et al., 2016).

Further, we investigated if there are any changes in neuronal activity due to HP in C57 and App mice. We observed dissociative effects of HP in resting state functional connectivity analysis, where HP leads to hyper-connectivity in C57 mice and hypo-connectivity in App mice. Interestingly, hyper-connectivity has been hypothesised to be a compensatory strategy against the progression of disease pathology/cognitive impairment (Carmichael et al., 2005; Palop et al., 2006; Di Filippo et al., 2008; Sigler et al., 2009; Mohajerani et al., 2011; Hillary et al., 2015; Siegel et al., 2016; Delli Pizzi et al., 2019). Although the components of this compensatory mechanism are unknown but during the disease progression, hyper-synchronous activity increases initially but decreases as the disease progresses, shifting to hypo-synchronous activity (Hillary and Grafman, 2017; Shah et al., 2018; Bing et al., 2019; Latif-Hernandez et al., 2019). A recent study suggests that cerebral remodeling during the early stage of modest cerebral hypoperfusion (as shown by increase in neuronal connectivity) could be in part mediated by paracrine interleukin 6 (IL-6) - a pro inflammatory cytokine, however, this may have detrimental effects in the long-term (Kuffner et al., 2022).

Analysis of sensory evoked cortical activations revealed alterations in sensory processing due to AD and HP. For stimulation of limbs ipsilateral to OH a significant reduction cortical activation was observed in both OH and NoH of C57-HP and App-HP mice, however, stimulation of limbs contralateral to OH led to an increased response in the NoH for C57-HP mice. These results suggest that for C57-HP mice, NoH might be compensating for the hypoperfusion in OH, however, for App-HP such a compensatory mechanism may not be plausible, probably due to underlying AD pathology. Interestingly, after targeted mini-strokes (Mohajerani et al., 2011) increased cortical activation was observed in the contralateral hemisphere and peri-infarct region on the stroke side for unaffected forelimb stimulation (ipsilateral to stroke hemisphere), suggesting that the network re-organization/response could be different for ischemic vs. chronic reduction of blood flow in the cortex.

We propose that increased inflammation due to HP could be leading to hyper-synchronous activity in C57-HP mice, but extensive inflammation in combination with Aβ plaque deposition in App-HP mice could be driving the change to the hypo-synchronous activity. Interestingly in mouse models of AD at pre-Aβ plaque deposition stage, increased microgliosis has been observed (Beauquis et al., 2014) in addition to hyper connectivity/activity, which changes to hypo-connectivity/activity as Aβ deposition and microgliosis increases (Shah et al., 2016; Shah et al., 2018; Latif-Hernandez et al., 2019). Microglia also interact with neurons in an activity-dependent manner after cerebral ischemia (Szalay et al., 2016), this may explain the increase in connectivity/activity associated with increased microgliosis in C57-HP mice. This also suggests that by increasing the burden on microglia and other glial cells, the brain can compensate for reduced blood flow and potentially other pathology (Venkat et al., 2016). Although some studies have shown that microglia are not necessary for Aβ plaque formation and maintenance, selective microglia removal is also shown to rescue memory functions and reduce disease pathology (Sosna et al., 2018; Kakae et al., 2019). Thus, further studies are needed to test the hypothesis that microglia activation could be a means to compensate for cognitive and functional dysfunctions associated with Aβ burden and hypoperfusion.

In conclusion, this study presents evidence of pathological and cortical dynamics alteration due to unilateral gradual cerebral hypoperfusion in C57BL/6J mice and a single App knock-in mouse model of AD (*App*^*NL-G-F*^). This gradual and mild form of cerebral HP mimics the AD risk factors such as hypercholesteremia, obesity, and atherosclerosis as these occur over a lifetime, gradually reducing blood flow to the brain, and do not have immediate onset. In contrast to our initial hypothesis that cerebral hypoperfusion may lead to Aβ plaque deposition, no Aβ plaques were observed in C57-HP mice although there was an increase in microgliosis, one reason for this could be the short duration of chronic HP, maybe if the duration of HP is extended one can expect to observe some Aβ pathology. However, in App-HP mice we do see an increase in Aβ plaques and microgliosis. Further, we found a dissociative effect of HP on cortical functional connectivity and sensory evoked activity, which, we hypothesize to be mediated in part by microglia. We propose that increased inflammation in the early stages of cerebral hypoperfusion can induce cerebral remodeling, however in the late stages or in the presence of AD related pathologies it may have detrimental effects. The results presented also lays an emphasis on how these risk factors may exacerbate AD pathology even though behavioural changes may not be immediately visible. Since identifying the impact of HP on AD pathology is not straightforward due to co-occurrence of other neuropathological features in humans such as neurofibrillary tangles, atrophy, inflammation, vascular amyloidosis, etc. (Austin et al., 2011), this study adds weight to the current literature on HP and AD as we were able to assess disease pathology, cognition and identify changes functional network connectivity/activity associated with HP/AD. Future experiments validating the potential mechanisms for the progression of AD and the role of vascular factors in cognitive decline and network dysfunctions are still necessary.

## Materials and Methods

All experimental procedures were approved by the institutional animal care committee and performed in accordance with the standards set out by the Canadian Council for Animal Care. Naïve male and female pairs of C57 (n = 19) and App (n = 27) mice bred in a pathogen free facility were used. UCAgO surgery was completed at two months of age, cerebral blood flow was measured following surgery, behavioural testing was done at 4 months, VSD imaging was completed following behavioural testing, and immunohistochemistry was completed at the end of all testing. UCAgO procedure and CBF measurements were described previously (Mehla, Lacoursiere, et al., 2018). Behavioural testing has been described previously (Mehla et al., 2018b; Mehla et al., 2018a; Mehla et al., 2019).

For *in vivo* VSD imaging mice were anesthetized with isoflurane (1.2–1.5%) for induction, followed by urethane for data collection (1.0-1.2 mg/kg, i.p). RH1691 dye (Optical Imaging, New York, NY) was applied to the cortex for 30-45 min. The voltage sensitive dye was excited with a red LED (Luxeon K2, 627 nm center), and excitation filters 630 ± 15 nm. Images were taken through a macroscope composed of front-to-front video lenses (8.6 × 8.6 mm field of view, 67 μm per pixel). The depth of field of the imaging setup used was ∼1 mm. To stimulate the forelimbs and hindlimbs, a 1 mA, 1-ms electrical pulse was delivered. The baseline of the optical signal (F_0_) was estimated, and the fluorescence changes were quantified as (F−F_0_)/F_0_ × 100%; F represents the fluorescence signal at any given time.

Aβ plaque was stained with 82E1 immunohistochemical markers. Microglial cells were stained with an ionized calcium-binding adapter molecule 1 (Iba1). A Nanozoomer serial slide scanner (NanoZoomer Digital Pathology 2.0-RS, HAMAMATSU, JAPAN) and Laser Scanning confocal microscope were used for imaging. Quantification of pathology was done using iLastik (Version 1.3.0-OSX) (Berg et al., 2019) and ImageJ software.

GraphPad Prism 7 for Mac OS X, v.7.0D (GraphPad Software, La Jolla California USA, www.graphpad.com) was used for statistical analysis of behavioral and pathology quantification, *p* value < 0.05 was considered statistically significant, adjusted *p* values reported. For spontaneous VSDI data a Generalized linear mixed-effects (GLME) model in MATLAB 2018b was used to predict correlation values with a fixed effect for group, including random effects for inter-regional correlations. Significance was set at α ≤ 0.05. Further, bootstrapping (resampling with replacement, 1000 samples) was used to determine 95% confidence intervals (CI) of condition mean differences (McGirr et al., 2017). Data is presented as mean ± SEM.

## SUPPLEMENTARY METHODS

### Animals and experimental timeline

Naïve male and female pairs of C57BL/6J (C57) and APP-KI mice carrying Arctic, Swedish, and Beyreuther/Iberian mutations (App) (25-30 g) bred in a pathogen free facility were used. The APP-KI mice were gifted by RIKEN Center for Brain Science, Japan. Mice were housed 4-5 mice per cage with *ad libitum* access to standard rodent chow and water and maintained on a 12-hour light/dark cycle. Colony room temperature was maintained at 21° C ± 1°. All experimentation was completed during the light cycle at the same time each day. All experimental procedures were approved by the institutional animal care committee and performed in accordance with the standards set out by the Canadian Council for Animal Care.

At two months of age, mice were randomly divided into sham or HP groups. The C57 sham (*n* = 13) and App sham (*n* = 11) groups consisted of mice that underwent sham surgery. The C57 HP (*n* = 6) and App HP (*n =* 16) groups were mice that underwent unilateral common carotid artery gradual occlusion (UCAgO). Behavioral testing started two months after surgery. Once behavioral testing was finished, VSDI imaging was done. At the experimental end point, mice were perfused, and tissue was collected for immunohistochemistry (Fig. 1).

### Unilateral common carotid artery gradual occlusion (UCAgO) surgery

The surgical procedure performed was described in a previous study (Mehla et al., 2018a). Briefly, mice were anesthetized with 1.5% isoflurane and a midline cervical incision exposed the common carotid artery (CCA), and the CCA was separated from the sheaths. The artery was lifted and placed in the internal lumen of the ameroid constrictor (AC, Research Instruments NW, 30094 Ingram Rd, Lebanon, OR 97355, USA; inner diameter, 0.5 mm; outer diameter, 3.25 mm, length, 1.28 mm) located just below the carotid bifurcation on the left side (Fig. 1B). The sham surgery followed the same protocol but without implanting an ameroid constrictor. The midline incision was sutured, and the mice were transferred to a recovery room.

### Laser Speckle flowmetry

Relative CBF was measured pre-surgery and on day 1, 3, 7, 14, & 28 post-surgery using laser speckle flowmetry, which has a linear relationship with absolute CBF values and obtains high spatial resolution 2D imaging as described in previous studies (Ayata et al., 2004; Mohajerani et al., 2011; Winship et al., 2014). The recordings were performed through a glass cover slip cranial window under anesthesia with 1.0 - 1.2% isoflurane (Mostany and Portera-Cailliau, 2008; Kyweriga et al., 2017). The mean CBF was measured was from identically sized ROI (located 2 mm lateral and 1 mm posterior from bregma) using ImageJ as described previously (Mohajerani et al., 2011; Winship et al., 2014). The reflectance optical signals reflect the CBF of the surface micro vessels in the cortex (Winship, 2014). CBF values are expressed as a percentage of the pre-surgery value. The subjects (*n* = 4) used for CBF were different from those used for behavioral assessment and histology in both sham and HP groups.

### Behavioural testing

Behavioural characterization was done at 4 months of age. Mice were handled for at least three days before starting the behavioral tasks. Spatial learning and memory were assessed using the Morris Water Task (MWT), Novel Object Recognition (NOR) was used to assess object learning and memory, and the Balance Beam (BB) test was performed to assess the sensory motor function.

MWT: Mice were trained on the MWT, as described previously in order to investigate spatial navigation learning and memory (Mehla et al., 2018b; Mehla et al., 2019). The acquisition phase consisted of four trials (60 sec maximum) per day for eight days. The trial was stopped once the mouse found the platform or 60 seconds elapsed, whichever occurred first. Mice were guided to the platform if they failed to find the platform. An intertrial interval of five minutes was used. Latency, pathlength, and swim speed were measured during the acquisition phase. On the ninth day, a single 60 sec no-platform probe trial was done. Mice were placed at a novel starting location opposing the target quadrant and allowed to swim freely for 60 sec before the trial ended. The percent of time spent in the target and non-target quadrants was measured during the no-platform probe trial.

NOR: The NOR was conducted to investigate object memory in mice as described previously (Mehla et al., 2018a). Mice were habituated to the testing box (White plastic, 52 × 51 × 30 cm; standard mouse bedding bottom) for five minutes for four days before testing. On the training day mice explored two identical objects for ten minutes. On the testing day, 24 hours later, a novel object replaced a familiar object, and mice explored for five minutes. Each trial started with a clean box and objects were cleaned with 70% isopropyl alcohol. Mice started each trial opposing the objects location. The investigation ratio (IR), the total time investigating object A divided by the sum of the time investigating Object A and Object B, was used to control for the individual differences investigating objects between mice. The IR was analyzed from recorded videos by an investigator, blinded to the groups.

Balance Beam: The BB is used to assess sensorimotor function (Mehla et al., 2018a). Mice were trained to traverse a 100 cm long, 1 cm diameter steel beam. Mice were trained incrementally starting from 10 cm, then 50 cm, and finally 100 cm. The training was complete once the mouse fully traversed the beam three times. Testing was done 24 hours later. The average time of three trials to traverse the beam was recorded. Falling would end the trial.

### VSD imaging

At five months of age, craniotomy for VSD imaging was performed as described previously (Mohajerani et al., 2010; Mohajerani et al., 2013; Kyweriga and Mohajerani, 2016). Mice were anesthetized with isoflurane (1.2–1.5%) for induction, followed by urethane for data collection (1.0-1.2 mg/kg, i.p). Mice were transferred on a metal plate that could be mounted onto the stage of the upright macroscope, and the skull was fastened to a steel plate. A tracheotomy was performed on mice to assist with breathing before starting the craniotomy. A 7×8 mm bilateral craniotomy (bregma 2.5 to −4.5 mm, lateral 0 to 4 mm) was made and the underlying dura was removed. Body temperature was maintained at 37 ± 0.2 °C degrees using a heating pad with a feedback thermistor.

For *in vivo* VSDI, RH1691 dye (Optical Imaging, New York, NY) was applied to the cortex for 30-45 min. For data collection, 12-bit images were captured with a CCD camera (1M60 Pantera, Dalsa, Waterloo, ON) and E8 frame grabber with XCAP 3.9 imaging software (EPIX, Inc., Buffalo Grove IL). The voltage sensitive dye was excited with a red LED (Luxeon K2, 627 nm center), and excitation filters 630 ± 15 nm (Mohajerani et al., 2010; Mohajerani et al., 2013; Chan et al., 2015). Images were taken through a macroscope composed of front-to-front video lenses (8.6 × 8.6 mm field of view, 67 μm per pixel). The depth of field of the imaging setup used was ∼1 mm (Lim et al., 2012). To stimulate the forelimbs and hindlimbs, thin acupuncture needles (0.14 mm) were inserted into the paws, and a 1 mA, 1-ms electrical pulse was delivered.

### VSD data analysis

VSD imaging of spontaneous cortical activity was recorded in the absence of visual, olfactory, tactile, or auditory stimulation during 15 min epochs with 10 ms (100 Hz) temporal resolution. Data was first denoised by applying singular-value decomposition and taking only the components with greatest associated singular values. The baseline of the optical signal (F_0_) captured from each pixel in the imaging window was calculated using the *locdetrend* function in the Choronux toolbox was used to fit a piecewise linear curve to the pixel time series using the local regression method (Mitra and Bokil, 2008). The fluorescence changes were quantified as (F−F_0_)/F_0_ × 100%; F represents the fluorescence signal at any given time and F_0_ represents the baseline of the optical signal. A band pass filter was applied (0.5–6 Hz) FIR filter on the Δ*F*/*F*_0_ signal as most of the optical signal power is concentrated in low frequencies (Mohajerani et al., 2013). Sensory stimulation was used to determine the coordinates for the primary hindlimb and forelimb sensory areas (HLS1 and FLS1). From these primary sensory coordinates, the relative locations of additional areas: parietal associational area (ptA), retrosplenial cortex (RS), medial secondary visual cortex (V2M), primary visual cortex (V1), lateral secondary visual cortex (V2L), barrel cortex (BCS1), hindlimb motor cortex (mHL), forelimb motor cortex (mFL), anterior segment of the secondary motor (aM2), and posterior segment of the secondary motor (pM2) were estimated using stereotaxic coordinates (Paxinos & Franklin, 2004). For region-based analyses, 24, 5 × 5-pixel ROIs were selected (12 from each hemisphere) in C57 sham, C57 HP, App sham, and App HP mice (*n* = 4, 4, 7, and 8 respectively). The regional functional connectivity matrix was generated using the zero-lag pearson correlation of ROI time courses.

VSD responses to sensory-evoked stimulation were calculated as the normalized difference to the average baseline estimated by fitting a fourth-degree polynomial (ΔF/F_0_ × 100) using custom-written code in MATLAB 2019b (Mathworks). Average sensory evoked response was calculated from 20 trials of stimulation with an inter-stimulus interval of 10 s. Alteration in evoked population responses were compared based on the following six parameters: rise time, fall time, inter-hemispheric delay, peak amplitude (Δ*F*/*F*_0_), area under the curve (AUC) and laterality index. Peak Δ*F*/*F*_0_ of the evoked responses is the maximum value from the onset of the stimulus to 200 ms thereafter, AUC is the area under Δ*F*/*F*_0_ – time curve of evoked response from the onset of the stimulus to 200 ms afterward, rise-time is the time interval in which the ROI signal rises from 10% to 90% of the peak evoked activation, fall-time is the time interval in which the ROI signal falls from 90% to 10% of the peak evoked activation, inter-hemispheric delay is the time difference of peak evoked activation in occluded and non-occluded hemispheres, and laterality index is defined as the ratio of difference and sum of peak hemispheric activations [(peak contralateral – peak ipsilateral) / (peak contralateral + peak ipsilateral)].

### Network analysis

Custom written MATLAB scripts in addition to modified version of Bioinformatics (Lim et al., 2015) and Brain Connectivity Toolbox (Rubinov and Sporns, 2010) were used to create a network diagram from the correlation matrices. Node size is proportional to the strength of the connections per node and edges represents connections that were greater (green) or less (red) than 10% of the control connections (Fig. 2B&E).

### Immunohistochemistry procedures and quantification

Mice were transcardially perfused with 1X phosphate buffer solution (PBS) followed by 4% paraformaldehyde (PFA). The brains were post-fixed in 4% PFA for 24 hours, followed by cryoprotection in a 30% sucrose solution with 0.02% sodium azide for at least three days before sectioning. Frozen brains were sectioned (40 µm) on a sliding microtome. The sections were stored in 1X PBS and 0.02% sodium azide at 4°C until processed.

To quantify Aβ plaque, the brain sections were stained with 82E1 immunohistochemical markers. Microgliosis was measured by staining microglial cells with an ionized calcium-binding adapter molecule 1 (Iba1) marker and the number of activated microglia (Iba1+) was quantified in the HPC and cortex of (see *Table 1. Key reagents and resources* for antibodies used). Sections were co-stained with DAPI (0.01 mg/ml; 140 ul/slides with cover slip). A Nanozoomer serial slide scanner (NanoZoomer Digital Pathology 2.0-RS, HAMAMATSU, JAPAN) and Laser Scanning confocal microscope were used for imaging. Quantification of pathology was done using iLastik (Version 1.3.0-OSX) (Berg et al., 2019) and ImageJ software. To quantify pathology, single channel images were used. Thresholding the channel of interest was done to apply consistency among all images to ensure training and prediction accuracy in iLastik. As iLastik uses several parameters for automated counting, predictions were not based solely on intensity of signal alone. Images were exported at 2.5x magnification and the regions of interests were isolated and copied into a 3000 × 3000-pixel window in ImageJ. The scale for each image was set by using the scale bar on the initial image which gave the number of pixels per millimeter; this value was used to determine the size of the plaque in millimeters. Images were processed with iLastik to identify Aβ plaques and activated microglia.

**Table 1.** Key reagents and resources.

| REAGENT or RESOURCE | SOURCE | IDENTIFIER |
| --- | --- | --- |
| **Antibodies** |  |  |
| 82E1 (β-Amyloid 1-16 clone; Anti-β-amyloid (N), mouse, 1:1000). | IBL, 10323 | |
| Iba1 (anti-Iba1, rabbit, 1:1000). | Wako, 019-19741 |  |
| ChAT (anti-Choline Acetyltransferase, monoclonal rabbit, 1:5000) | Abcam, ab178850 |  |
| anti-mouse-Alexafluor-488 (goat, 1:1000) | Abcam, ab150113 |  |
| anti-rabbit-Alexafluor-594 (goat, 1:1000). | Invitrogen, A11037 |  |
| DAPI (0.01 mg/ml; 140 ul/slides) |  |  |
| Vectashield (H-1000 or H-1200) | Vector Labs |  |
| **Chemicals, Peptides, and Recombinant Proteins** |  |  |
| VSDI, RH1691 dye | Optical Imaging, New York, NY |  |
| **Experimental Models: Organisms/Strains** |  |  |
| App mice; homozygous; Male | RIKEN breeding pair (Saito et al., 2014); bred in house. |  |
| C57BL/6J; Male. | Bred in house. |  |
| **Software and Algorithms** |  |  |
| EPIX E8 frame grabber with XCAP 3.9 imaging software | EPIX, Inc., Buffalo Grove IL |  |
| Chronux toolbox | http://chronux.org/ |  |
| MATLAB 2018b | www.mathworks.com/ |  |
| GraphPad Prism 7 for Mac OS X, v.(7.0D) | GraphPad Software, La Jolla California USA,<br>www.graphpad.com |  |
| Brain Connectivity Toolbox | (Rubinov & Sporns, 2010) |  |
| NDP.view 2 |  |  |
| iLastik (Version 1.3.0-OSX) | (Berg et al., 2019) |  |
| ImageJ |  |  |
| **Other** |  |  |
| 1M60 Pantera, CCD camera | Dalsa, Waterloo, ON |  |
| Red LED (Luxeon K2, 627 nm center) |  |  |
| 673–703 nm bandpass optical filter | Semrock, New York, NY |  |
| Ameroid constrictor; inner diameter, 0.5 mm; outer diameter, 3.25 mm, length, 1.28 mm | Research Instruments NW, 30094 Ingram Rd, Lebanon, OR 97355, USA |  |
| NanoZoomer Digital Pathology 2.0-RS | HAMAMATSU, JAPAN |  |

### Statistical Analysis

GraphPad Prism 7 for Mac OS X, v.7.0D (GraphPad Software, La Jolla California USA, www.graphpad.com) was used for statistical analysis of behavioral and pathology quantification. A *p* value < 0.05 was considered statistically significant, adjusted *p* values reported. A two-way repeated measures ANOVA followed by Sidak’s multiple comparison was used to determine significance between CBF across days and between the occluded hemisphere and non-occluded hemispheres (Fig. 1C). For cortical evoked activations 3-way ANOVA with Bonferroni post-hoc correction (α = 0.01), was used to study the effect of regions, HP and strain. Effects of STRAIN, HP, and DAY were assessed with 3-way ANOVA and Tukey’s multiple comparison significance in behavioural experiments; sphericity was corrected with the Giesser-Greenhouse correction (Fig. 6). For spontaneous VSDI data a Generalized linear mixed-effects (GLME) model in MATLAB 2018b was used to predict correlation values with a fixed effect for group, including random effects for inter-regional correlations. Significance was set at α ≤ 0.05. Further, bootstrapping (resampling with replacement, 1000 samples) was used to determine 95% confidence intervals (CI) of condition mean differences (McGirr et al., 2017) (Fig. 2). An ordinary one-way ANOVA was used to analyze the groups and between ipsilateral and contralateral sides and between groups for Aβ and microgliosis pathology. Region specific effects shown in fig. S2, S3, S6 & S7 were calculate with two-tailed two-sample *t*-test. Results presented as mean ± standard error of the mean (SEM).

## Acknowledgments

This work was supported by Natural Sciences and Engineering Research Council of Canada, Canada Discovery Grant #40352, #06347, and #03857 to MHM, RJM, and RJS, respectively, Alberta Innovates (MHM), Alberta Alzheimer Research Program (MHM, RJS, RJM), Alzheimer Society of Canada (MHM), Alberta Prion Research Institute (MHM and RJS), and Canadian Institute for Health Research (MHM, RJM, and RJS). The authors thank Dr Takashi Saito and Prof. Takaomi C Saido from “Laboratory for Proteolytic Neuroscience RIKEN Center for Brain Science, Wako-shi, Saitama, Japan” for providing the *App*^*NL-G-F*^ mice as a gift. The authors also thank Di Shao and Behroo Mirza Agha for animal breeding, Jianjun Sun for his surgical expertise for the VSD imaging craniotomy procedures and Valerie Lapointe for immunohistochemistry. We would also like to thank the University of Lethbridge Animal Care Staff.

## Author Contributions

Conceptualization, M.H.M., S.G.L., S.S. J.M. and R.J.M.; Experimentation, J.M. (UCAgO surgeries, behavior, immunohistochemistry, and CBF imaging), S.G.L. (behavior and immunohistochemistry), M.N. and S.S. (optical imaging), Formal Analysis, J.M. (CBF analysis), S.G.L. (behavior, immunohistochemistry), and S.S. (optical imaging); Writing - Original Draft, S.S., S.G.L., and J.M.; Writing - Review & Editing, M.H.M., S.G.L., S.S., M.N., J.M. R.J.M., and R.J.S.; Funding Acquisition, M.H.M., R.J.M., and R.J.S.; Supervision, M.H.M., R.J.M., and R.J.S.

## Data Availability

The dataset generated in the current study will be made available by the corresponding authors on reasonable request.

## Figure Legends

**Figure S1.**
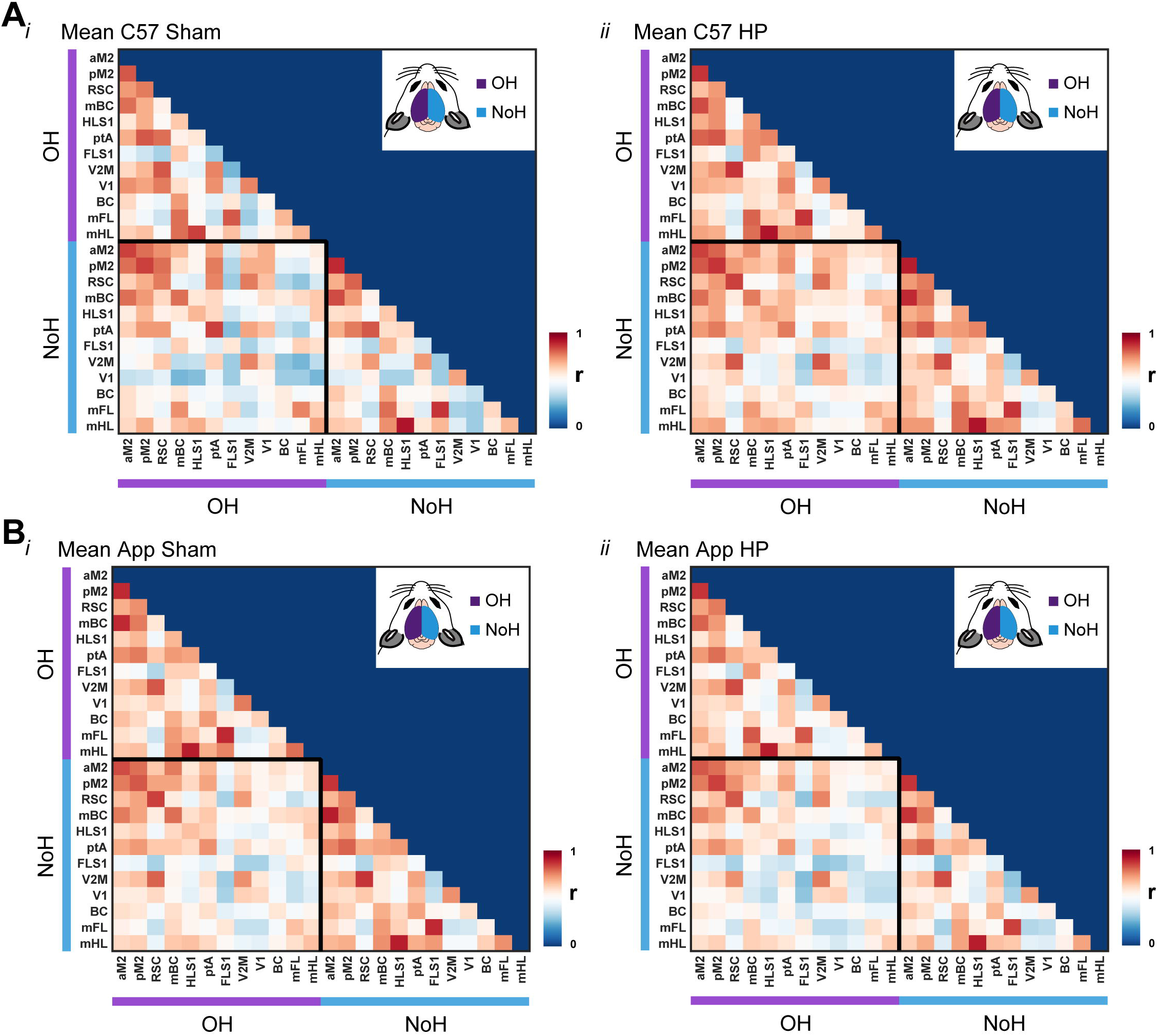
Cortical correlation matrix. Mean of cortical correlation matrix of (A*i-ii*) C57 mice (*n* = 4), and C57 HP mice (*n* = 4). (B*i-ii*) App mice (*n* = 7), and App HP mice (*n* = 8).

**Figure S2.**
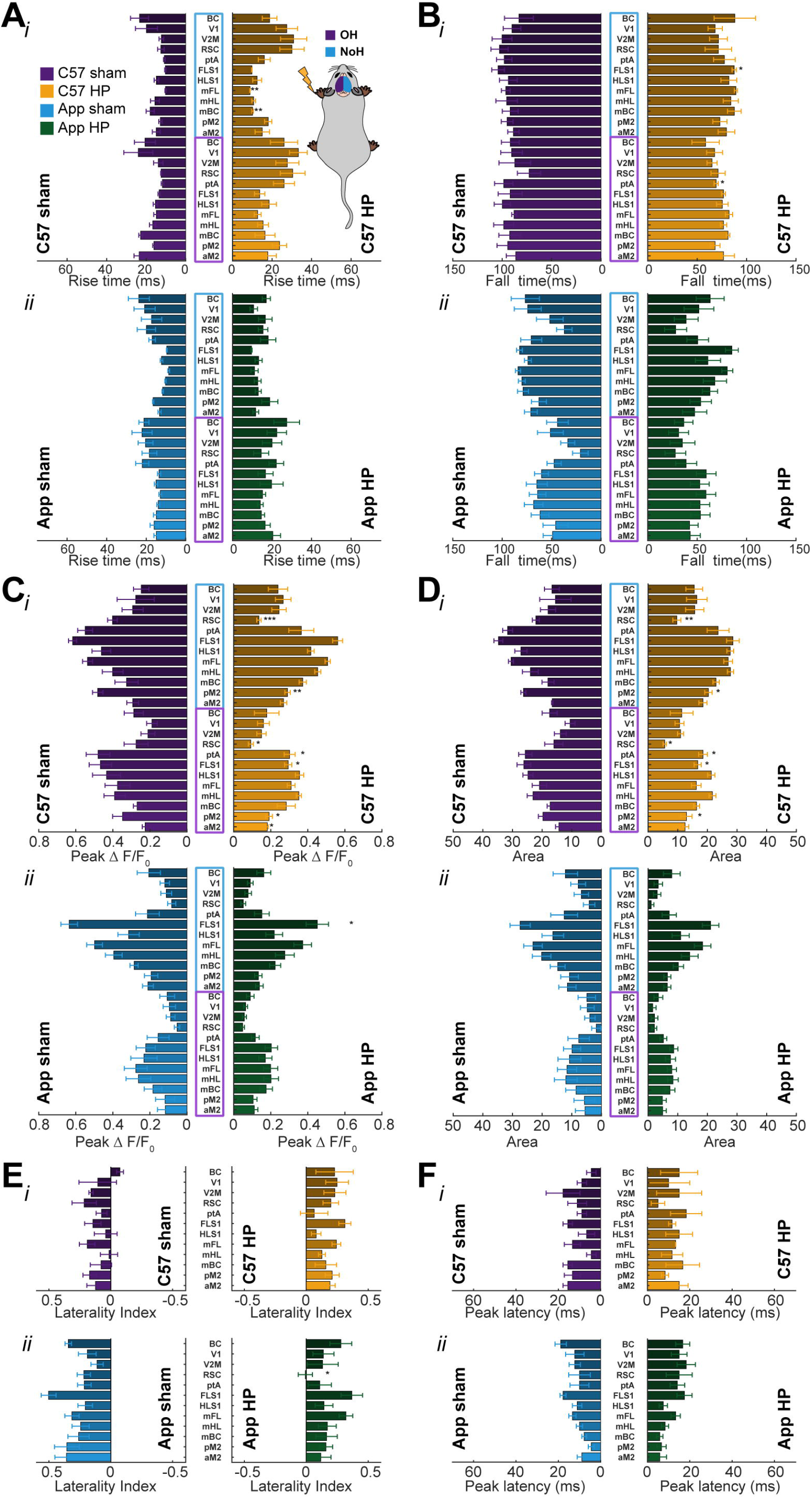
Region specific changes for left forelimb (FL) stimulus evoked cortical activations. Region specific comparison of (A*i-ii*) rise time (10% to 90% of peak), (B*i-ii*) fall time (90% to 10% of peak), (C*i-ii*) peak change in fluorescence (Δ*F*/*F*_0_), (D*i-ii*) area under Δ*F*/*F*_0_ – time curve (AREA), (E*i-ii*) laterality index [(peak contralateral – peak ipsilateral) / (peak contralateral + peak ipsilateral)], and (F*i-ii*) peak latency [abs (peak time OH – peak time NoH)], in C57 and App mice. Each parameter was extracted from evoked cortical activations in twenty-four (24), 5 × 5-pixel regions of interest (ROIs) (12 from each hemisphere). Primary hindlimb and forelimb sensory areas (HLS1 and FLS1), parietal associational area (ptA), retrosplenial cortex (RS), medial secondary visual cortex (V2M), primary visual cortex (V1), lateral secondary visual cortex (V2L), barrel cortex (BCS1), hindlimb motor cortex (mHL), forelimb motor cortex (mFL), anterior segment of the secondary motor (aM2), and posterior segment of the secondary motor (pM2), as estimated using stereotaxic coordinates (Paxinos & Franklin, 2004). The purple outline around region names along y-axis represents ROIs in OH and the blue represents ROIs in NoH. App animals (n = 7 Sham; n = 7 HP) and C57 animals (n = 4 Sham; n = 4 HP). All values are expressed as mean ± SEM. * *p* < 0.05; ** *p* < 0.01; *** *p* < 0.001.

**Figure S3.**
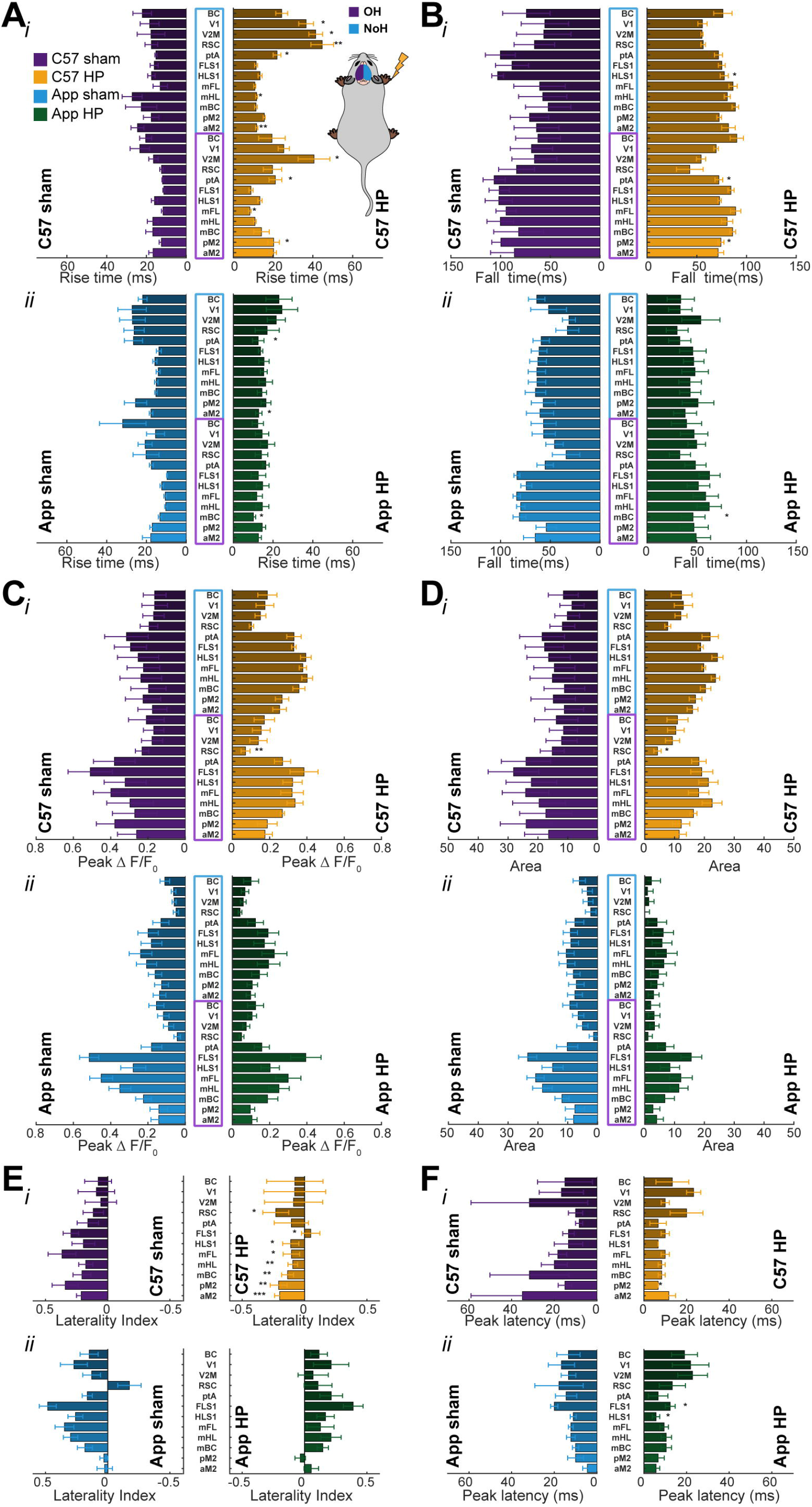
Region specific changes for right forelimb (FL) stimulus evoked cortical activations. Region specific comparison of (A*i-ii*) rise time (10% to 90% of peak), (B*i-ii*) fall time (90% to 10% of peak), (C*i-ii*) peak change in fluorescence (Δ*F*/*F*_0_), (D*i-ii*) area under Δ*F*/*F*_0_ – time curve (AREA), (E*i-ii*) laterality index [(peak contralateral – peak ipsilateral) / (peak contralateral + peak ipsilateral)], and (F*i-ii*) peak latency [abs (peak time OH – peak time NoH)], in C57 and App mice. Each parameter was extracted from evoked cortical activations in twenty-four (24), 5 × 5-pixel regions of interest (ROIs) (12 from each hemisphere; see Methods). The purple outline around region names along y-axis represents ROIs in OH and the blue represents ROIs in NoH. App animals (n = 7 Sham; n = 7 HP) and C57 animals (n = 4 Sham; n = 4 HP). All values are expressed as mean ± SEM. * *p* < 0.05; ** *p* < 0.01; *** *p* < 0.001.

**Figure S4.**
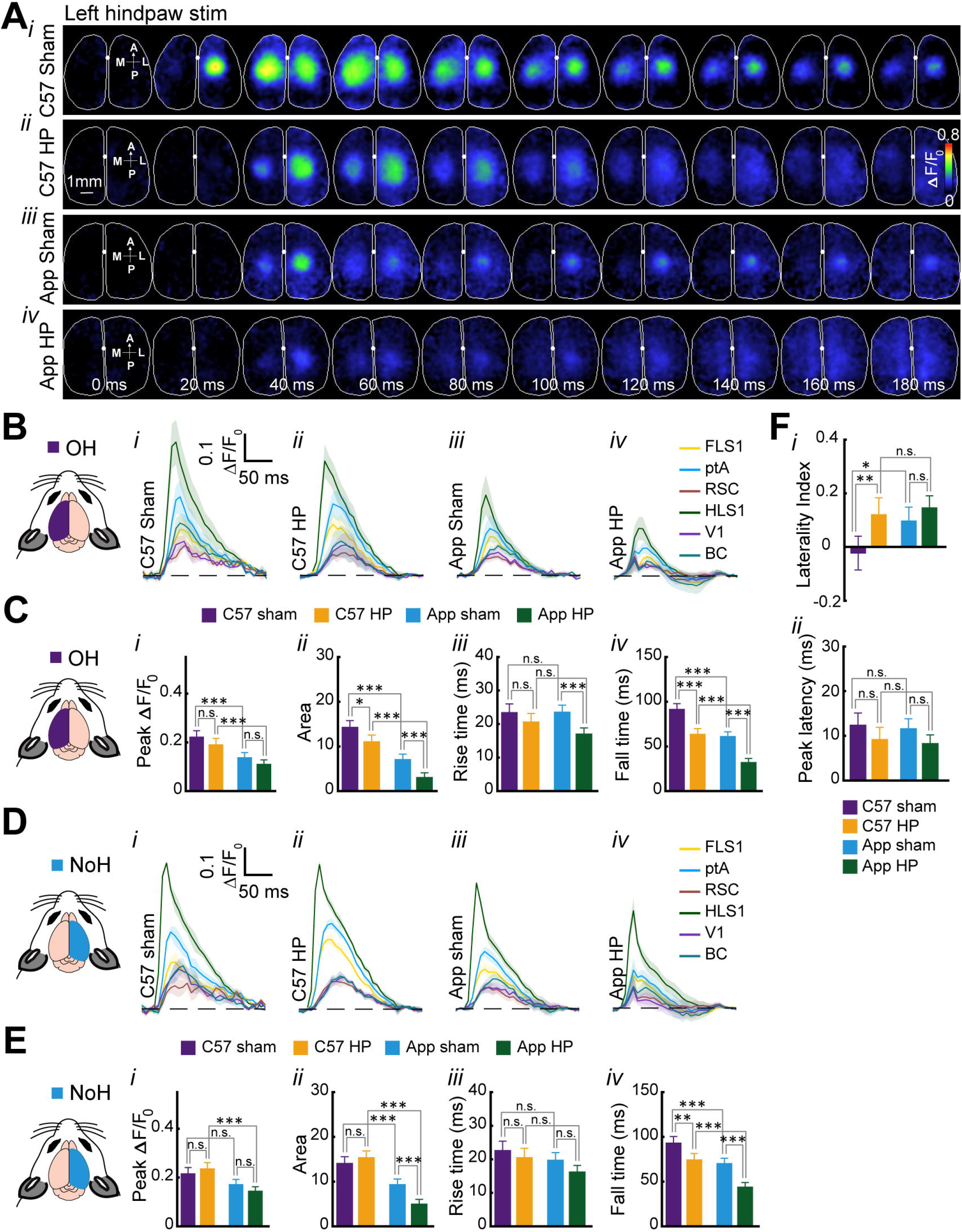
Reduced left hindlimb (HL) stimulus evoked cortical activations due to HP in C57 and APP mice. (A*i-iv*) Representative patterns of bilateral cortical activation following 1 mA, 1 ms pulse stimulation to left HL of C57 sham, C57 HP, App sham, and App HP mice. The VSDI montages represents 10 frames of evoked cortical responses at interval of 20 ms after stimulus onset (0.00 ms). The first image in the montage indicates the anterior (A), posterior (P), medial (M) and lateral (L) directions. (B*i-iv*,D*i-iv*) Average VSD signal from representative six regions of interest (ROIs) of OH and NoH in response to left hindlimb stimulation. Primary hindlimb and forelimb sensory areas (HLS1 and FLS1), parietal associational area (ptA), retrosplenial cortex (RS), primary visual cortex (V1), barrel cortex (BC), as estimated using stereotaxic coordinates (Paxinos & Franklin, 2004). (C) In OH, there was a significant reduction in area under Δ*F*/*F*_0_ – time curve (*p* < 0.0001) (C*ii*), rise time (C*iii*) (*p* < 0.0001) and fall time (C*iv*) (*p* < 0.001) for App HP group compared to App sham group. Similar differences in area under Δ*F*/*F*_0_ – time curve (*p* < 0.05) and fall time (*p* < 0.0001) were also observed for C57 HP and C57 sham group, however, no significant changes were observed in peak Δ*F*/*F*_0_ (C*i*) across C57/App sham and HP groups. (E) In NoH, significant reduction area under Δ*F*/*F*_0_ – time curve (E*ii*) (*p* < 0.0001) and fall time (E*iv*) (*p* < 0.0001) was observed for App sham and App HP group, but there was significant decrease in fall time for C57 HP group as compared to C57 sham group (*p* < 0.01). Further, no significant changes were observed in peak Δ*F*/*F*_0_ (E*i*) across C57/App sham and HP groups, and there was no significant difference in rise time (E*iii*) for any group. These results suggest reduced cortical activations in OH and NoH due to both HP and AD pathology for HL left stimulation. (F) Comparison of laterality index (F*i*) revealed significant increase in C57 HP group compared to C57 sham group (*p* < 0.01), but no difference was observed in peak latency (F*ii*) for any group. App animals (n = 7 Sham; n = 7 HP) and C57 animals (n = 4 Sham; n = 4 HP). All values are expressed as mean ± SEM. * *p* < 0.05; ** *p* < 0.01; *** *p* < 0.001.

**Figure S5.**
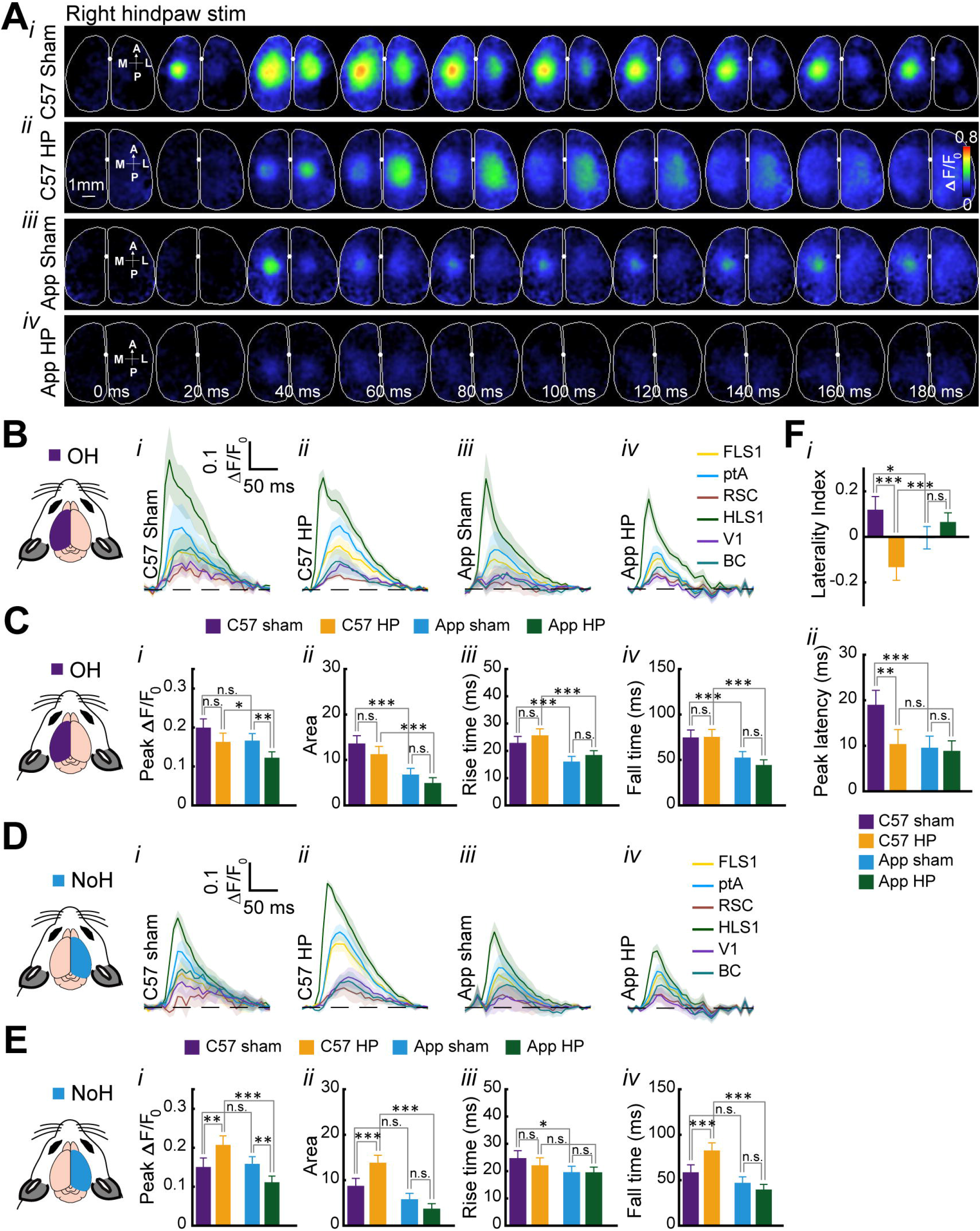
Increased right hindlimb (HL) stimulus evoked cortical activations in NoH of C57 HP mice. (A*i-iv*) Representative patterns of bilateral cortical activation following 1 mA, 1 ms pulse stimulation to right HL of C57 sham, C57 HP, App sham, and App HP mice. The VSDI montages represents 10 frames of evoked cortical responses at interval of 20 ms after stimulus onset (0.00 ms). The first image in the montage indicates the anterior (A), posterior (P), medial (M) and lateral (L) directions. (B*i-iv*, D*i-iv*) Average VSD signal from representative six regions of interest (ROIs) of OH and NoH in response to right hindlimb stimulation. Primary hindlimb and forelimb sensory areas (HLS1 and FLS1), parietal associational area (ptA), retrosplenial cortex (RS), primary visual cortex (V1), barrel cortex (BC), as estimated using stereotaxic coordinates (Paxinos & Franklin, 2004). (C) In OH, there was a significant reduction in peak Δ*F*/*F*_0_ (C*i*) (*p* < 0.01) in App HP mice compare to App sham mice, however no changes were observed in area under Δ*F*/*F*_0_ – time curve (C*ii*), rise time (C*iii*) and fall time (C*iv*) across C57/App sham and HP groups. (E) In NoH, there was significant increase in peak Δ*F*/*F*_0_ (E*i*) (*p* < 0.01), area under Δ*F*/*F*_0_ – time curve (E*ii*) (*p* < 0.0001) and fall time (E*iv*) (*p* < 0.001) of C57 HP mice as compared to C57 sham mice, however, there was significant decrease in peak Δ*F*/*F*_0_ of App HP group as compared to App sham group (*p* < 0.01). These results suggest reduced cortical activations in OH and NoH due to HP in App mice for HL right stimulation, however there was an increased cortical activation in NoH due to HP in C57 group. (F) Significant negative laterality index (F*i*) for C57 HP group (*p* < 0.0001) suggest that there is not only reduction of cortical activation in OH but also increase in NoH cortical activation, further a significant reduction in peak latency was observed only for C57 HP group as compared to C57 sham group (F*ii*) (*p* < 0.01). App animals (n = 7 Sham; n = 8 HP) and C57 animals (n = 4 Sham; n = 4 HP). All values are expressed as mean ± SEM. * *p* < 0.05; ** *p* < 0.01; *** *p* < 0.001.

**Figure S6.**
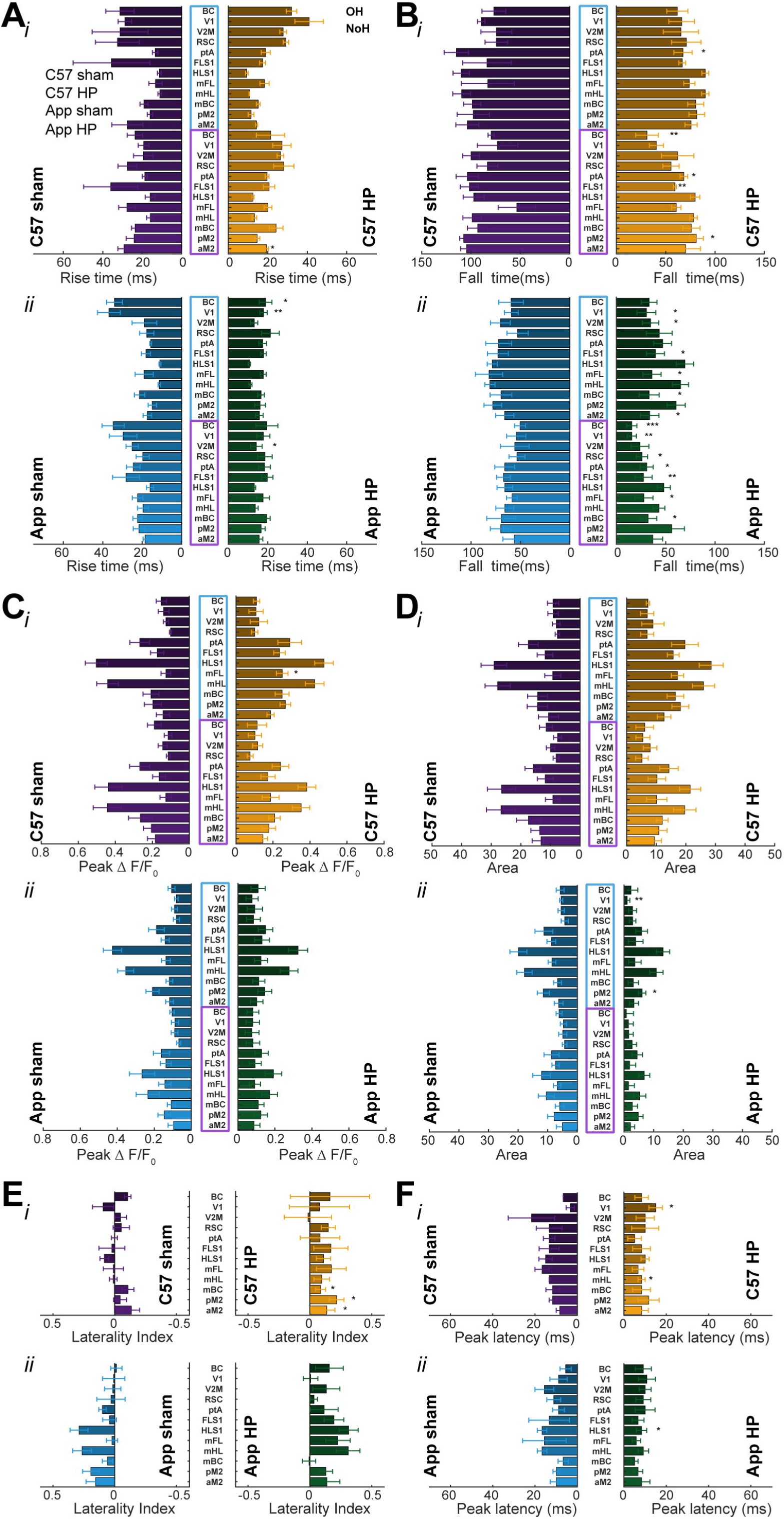
Region specific changes for left forelimb (HL) stimulus evoked cortical activations. Region specific comparison of (A*i-ii*) rise time (10% to 90% of peak), (B*i-ii*) fall time (90% to 10% of peak), (C*i-ii*) peak change in fluorescence (Δ*F*/*F*_0_), (D*i-ii*) area under Δ*F*/*F*_0_ – time curve (AREA), (E*i-ii*) laterality index [(peak contralateral – peak ipsilateral) / (peak contralateral + peak ipsilateral)], and (F*i-ii*) peak latency [abs (peak time OH – peak time NoH)], in C57 and App mice. Each parameter was extracted from evoked cortical activations in twenty-four (24), 5 × 5-pixel regions of interest (ROIs) (12 from each hemisphere; see Methods). The purple outline around region names along y-axis represents ROIs in OH and the blue represents ROIs in NoH. App animals (n = 7 Sham; n = 8 HP) and C57 animals (n = 4 Sham; n = 4 HP). All values are expressed as mean ± SEM. * *p* < 0.05; ** *p* < 0.01; *** *p* < 0.001.

**Figure S7.**
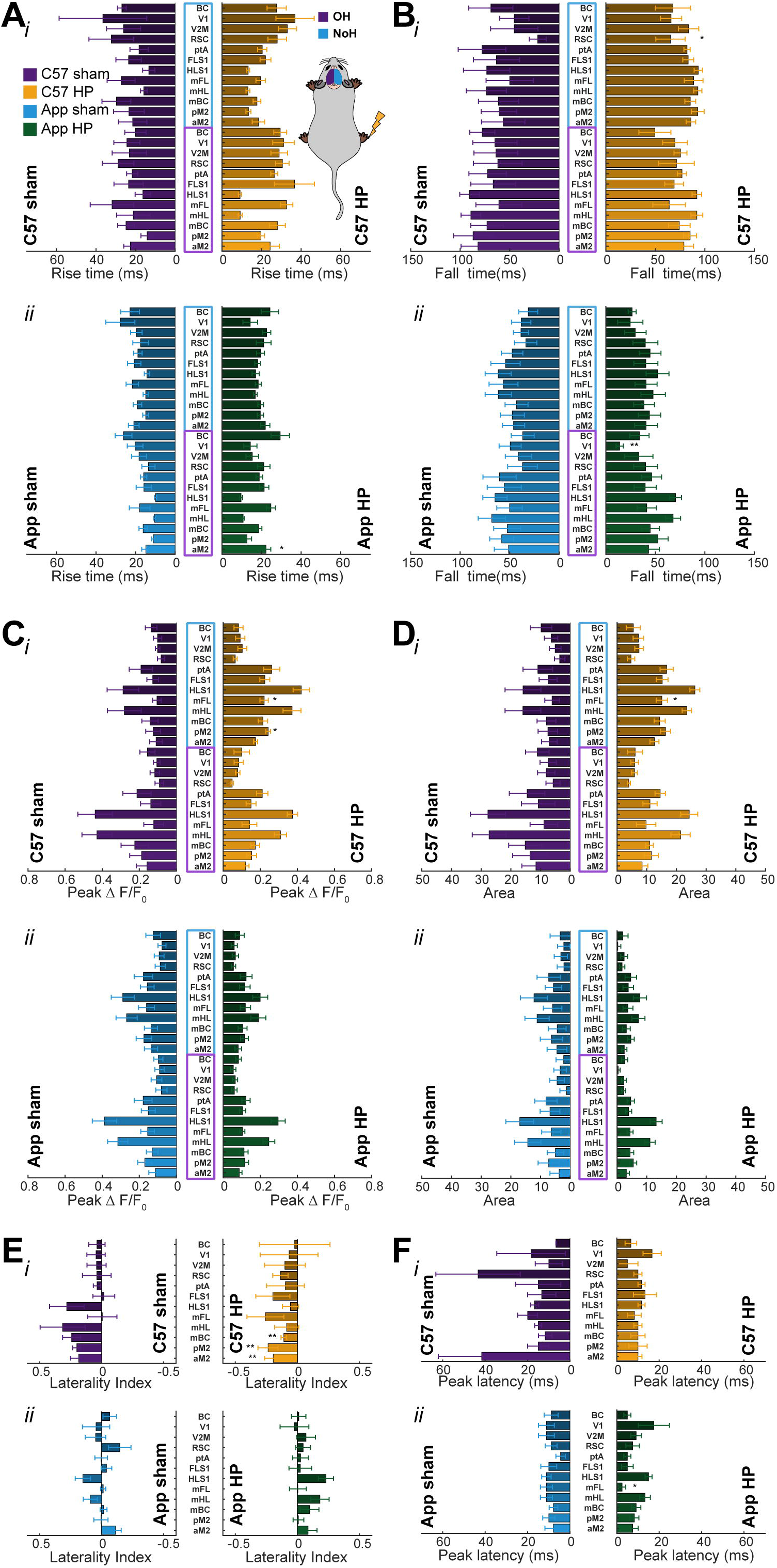
Region specific changes for right forelimb (HL) stimulus evoked cortical activations. Region specific comparison of (A*i-ii*) rise time (10% to 90% of peak), (B*i-ii*) fall time (90% to 10% of peak), (C*i-ii*) peak change in fluorescence (Δ*F*/*F*_0_), (D*i-ii*) area under Δ*F*/*F*_0_ – time curve (AREA), (E*i-ii*) laterality index [(peak contralateral – peak ipsilateral) / (peak contralateral + peak ipsilateral)], and (F*i-ii*) peak latency [abs (peak time OH – peak time NoH)], in C57 and App mice. Each parameter was extracted from evoked cortical activations in twenty-four (24), 5 × 5-pixel regions of interest (ROIs) (12 from each hemisphere; see Methods). The purple outline around region names along y-axis represents ROIs in OH and the blue represents ROIs in NoH. App animals (n = 7 Sham; n = 8 HP) and C57 animals (n = 4 Sham; n = 4 HP). All values are expressed as mean ± SEM. * *p* < 0.05; ** *p* < 0.01; *** *p* < 0.001.

**Figure S8.**
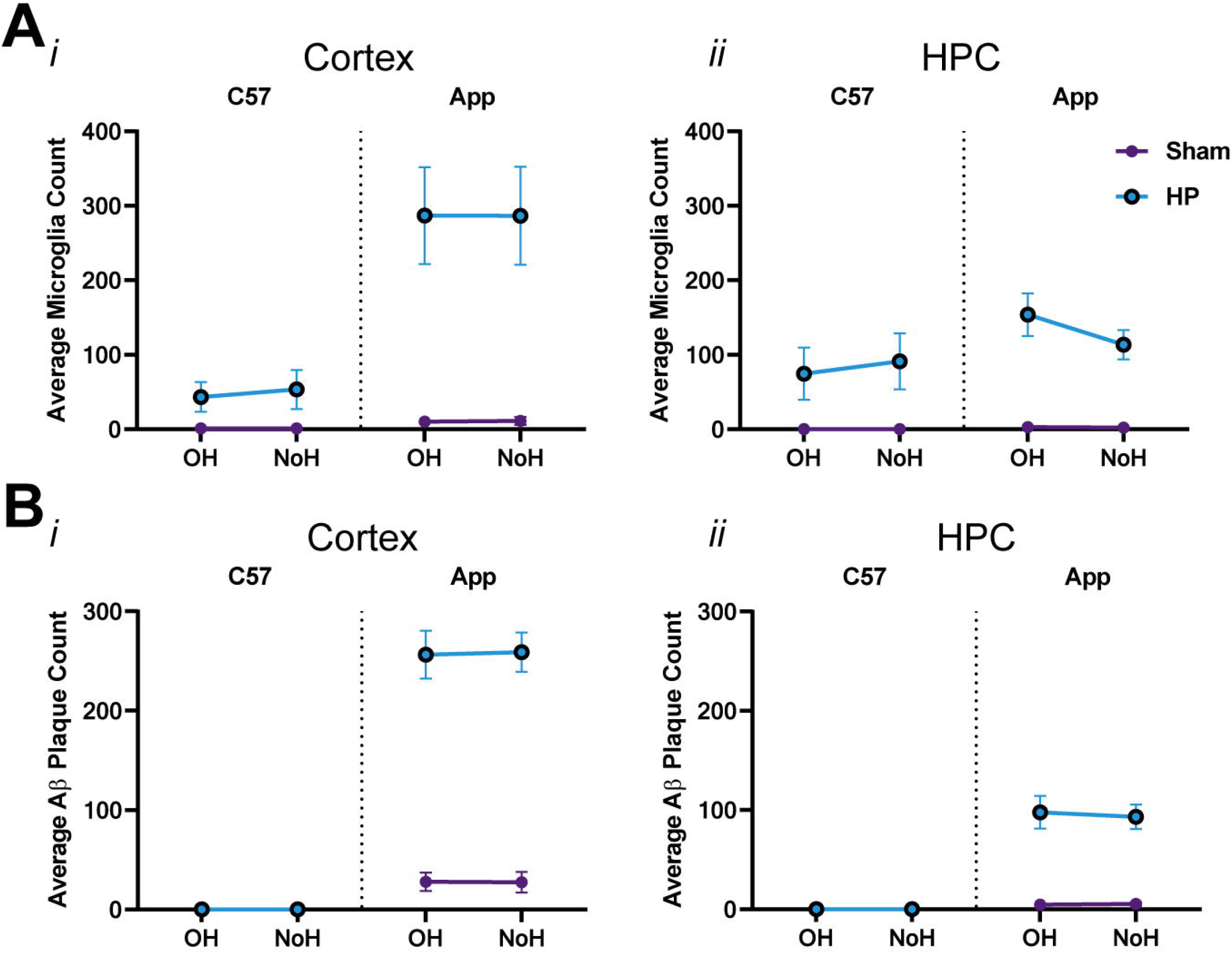
No hemisphere specific changes in microgliosis and Aβ pathology after HP. (A*i-ii*) activated microglial count in OH and NoH of Sham and HP mice. No hemisphere specific changes were observed in both cortex and HPC. (B*i-ii*) Similarly no hemisphere specific changes were observed in Aβ plaque count. * *p* < 0.05; ** *p* < 0.01; *** *p* < 0.001

